# Measuring the spatial heterogeneity on the reduction of vaginal fistula burden in Ethiopia between 2005 and 2016

**DOI:** 10.1101/2019.12.21.885632

**Authors:** Kebede Deribe, Claudio Fronterre, Tariku Dejene, Sibhatu Biadgilign, Amare Deribew, Muna Abdullah, Jorge Cano

## Abstract

Vaginal fistula is a shattering maternal complication characterized by an anomalous opening between the bladder and/or rectum and vagina resulting in continuous leakage of urine or stool. Although prevalent in Ethiopia, its magnitude and distribution is not well studied. We used statistical mapping models using 2005 and 2016 Ethiopia Demographic Health Surveys data combined with a suite of potential risk factors to estimate the burden of vaginal fistula among women of childbearing age. The estimated number of women of childbearing age with lifetime and untreated vaginal fistula in 2016 were 72,533 (95% CI 38,235–124,103) and 31,961 (95% CI 11,596–70,309) respectively. These figures show reduction from the 2005 estimates: 98,098 (95% CI 49,819–170,737) lifetime and 59,114 (95% CI 26,580–118,158) untreated cases of vaginal fistula. The number of districts having more than 200 untreated cases declined drastically from 54 in 2005 to 6 in 2016. Our results show a significant subnational variation in the burden of vaginal fistula. Overall, between 2005 and 2016 there was substantial reduction in the prevalence of vaginal fistula in Ethiopia. Our results help guide local level tracking, planning, spatial targeting of resources and implementation of interventions against vaginal fistula.

## Introduction

Vaginal fistula (VF), which is categorised as vesicovaginal or rectovaginal fistula, occurs when there is abnormal opening (fistula) between the bladder and/or the rectum and the vagina ^1^. It is most common in developing counties and often occurs after protracted obstructed labour, hence it is referred as obstetric fistula. The condition is highly debilitating and stigmatizing; women suffering from vaginal fistula are often stigmatized ^2–4^. VF is a disease deeply embedded in poverty, harmful practices such as early marriage and inequalities in access to health care. Obstetric fistula is virtually unheard of in developed countries.

Campaign to End Obstetric Fistula was launched in 2003 by The United Nations Population Fund and its implementing partners that sets an agenda for ‘the elimination of obstetric fistula’ in high burden countries^5^. In line with this global commitment, Ethiopia have targeted to eliminate obstetric fistula by 2020 ^6^. Data for VF are available at the national level in Africa but there is little information about subnational variations ^1,7,8^. Although a few studies have reported prevalence of vaginal fistula at specific sites ^1,7,9,10^, no studies have examined vaginal fistula prevalence at fine spatial resolution for entire countries. Analysis at country level, although useful for advocacy, may mask subnational geographical variation, which is critical for subnational level planning and measurement of inequalities in a country.

Some studies have been carried out at sub-national level of Ethiopia to establish the burden of obstetric fistula (OF). A study conducted in East Hararghe, South Gondar and West Gojjam over a randomly selected sample of 23,023 women of childbearing age (WCA) (15-49 years) sampled from 113 villages determined the prevalence of OF to be 0.6 per 1,000, of untreated fistula 0.2 per 1,000 and of symptomatic pelvic organ prolapse 10 per 1,000 of WCA ^9^. Another study included a total of 19,153 households from seven regions of Ethiopia. Untreated fistula prevalence was estimated at 1.5 per 1,000 WCA. The authors extrapolated their findings and roughly estimated that there were 26,819 women with untreated fistula in Ethiopia ^10^. Finally, another study based on data collected through Demographic and Health (DHS) surveys in 2005 estimated the life time and point prevalence of vaginal fistula to be 7.1 (5.5-8.7) and 5·6 (4·3−7·0) per 1,000 WCA, respectively ^7^. Although all the above studies contributed to the body of literature, none of them comprehensively reported local distribution and burden of fistula in the country. In addition, the statistics presented by these studies were based on raw collected data and did not account either for potential risk factors or for ongoing interventions, which have been conducted over the last decade and may have contributed to reduce the burden of vaginal fistula in Ethiopia.

The health system and health services in Ethiopia are devolved to regions and district level (*woredas*). Therefore, reliable data on the prevalence and burden of OF at district level is imperative for planning and action. However, such kind of data is lacking in Ethiopia despite the clear socio-economic difference in differ geographies ^7,8^. We aimed to assess the prevalence and burden of VF at subnational level and understand the geographical inequalities in the burden of VF. The 2005 and 2016 Ethiopian DHS data were extracted to estimate the prevalence and burden of VF in Ethiopia and trend in its distribution over time. Using georeferenced survey data and continuous surfaces of potential social determinants of vaginal fistula, we intended to provide estimates of vaginal fistula prevalence at a resolution as much as 1 square-km.

## Results

### General description

Ethiopian DHS for 2005 and 2016 collected data from 14,070 and 15,683 WCA, respectively. Table 1 shows the number of women with vaginal fistula symptoms and the number who have not been treated for this condition by region. There was no significant difference in the median of prevalence of untreated fistula between rural and urban communities across regions neither in 2005 (Wilcoxon’s test, p-value = 0.7545) nor in 2016 (Wilcoxon’s test, p-value = 0.1843). Neither was there significant difference in the median of prevalence of lifetime fistula between rural and urban communities in 2005 (Wilcoxon’s test, p-value = 0.439), although prevalence was found to be higher in rural than urban areas in 2016 (Wilcoxon’s test, p-value < 0.05). When we compared the prevalence of lifetime fistula in rural and urban areas between years, we observed no difference in the prevalence in rural areas (Wilcoxon’s test, p-value = 0.3381) whereas the prevalence in urban areas declined sharply from 2005 to 2016 (Wilcoxon’s test, p-value < 0.05). However, the prevalence of untreated fistula declined significantly both in rural and urban areas (Wilcoxon’s test, p-value <0.05) in 2016.

**Table 1.** Number of women of childbearing age surveyed in DHS 2005 and DHS 2016

| Region | 2005 |  |  | 2016 |  |  |
| --- | --- | --- | --- | --- | --- | --- |
|  | Total interviewed (n) | Women reporting vaginal fistula symptoms (n) | Untreated cases of vaginal fistula | Total interviewed (n) | Women reporting vaginal fistula symptoms (n) | Untreated cases of vaginal fistula |
| Tigray | 1257 | 14 | 10 | 1682 | 19 | 4 |
| Afar | 789 | 5 | 3 | 1128 | 6 | 1 |
| Amara | 1943 | 8 | 1 | 1719 | 12 | 9 |
| Oromia | 2230 | 20 | 17 | 1892 | 4 | 1 |
| Somali | 669 | 1 | 1 | 1391 | 5 | 1 |
| Benishangul Gumuz | 846 | 4 | 3 | 1126 | 7 | 4 |
| SNNP | 2087 | 26 | 19 | 1849 | 6 | 2 |
| Gambella | 729 | 5 | 1 | 1035 | 3 | 2 |
| Harari | 844 | 1 | 1 | 906 | 0 | 0 |
| Addis Ababa | 1869 | 14 | 7 | 1824 | 7 | 4 |
| Dire Dawa | 807 | 5 | 4 | 1131 | 1 | 0 |
|  | 14070 | 103 | 67 | 15683 | 70 | 28 |

### Estimated burden of vaginal fistula

The Fig 1 & 2 shows a heterogeneous spatial distribution and number of cases of lifetime and untreated VF in Ethiopia. The highest prevalence of VF for 2005 is estimated in some parts of Amhara, Oromia, Southern Nations, Nationalities, and Peoples (SNNP) and Tigray regions. The predicted map showed a significant reduction of lifetime and untreated fistula in 2016, although this reduction is spatially heterogeneous.

**Figure 1.**
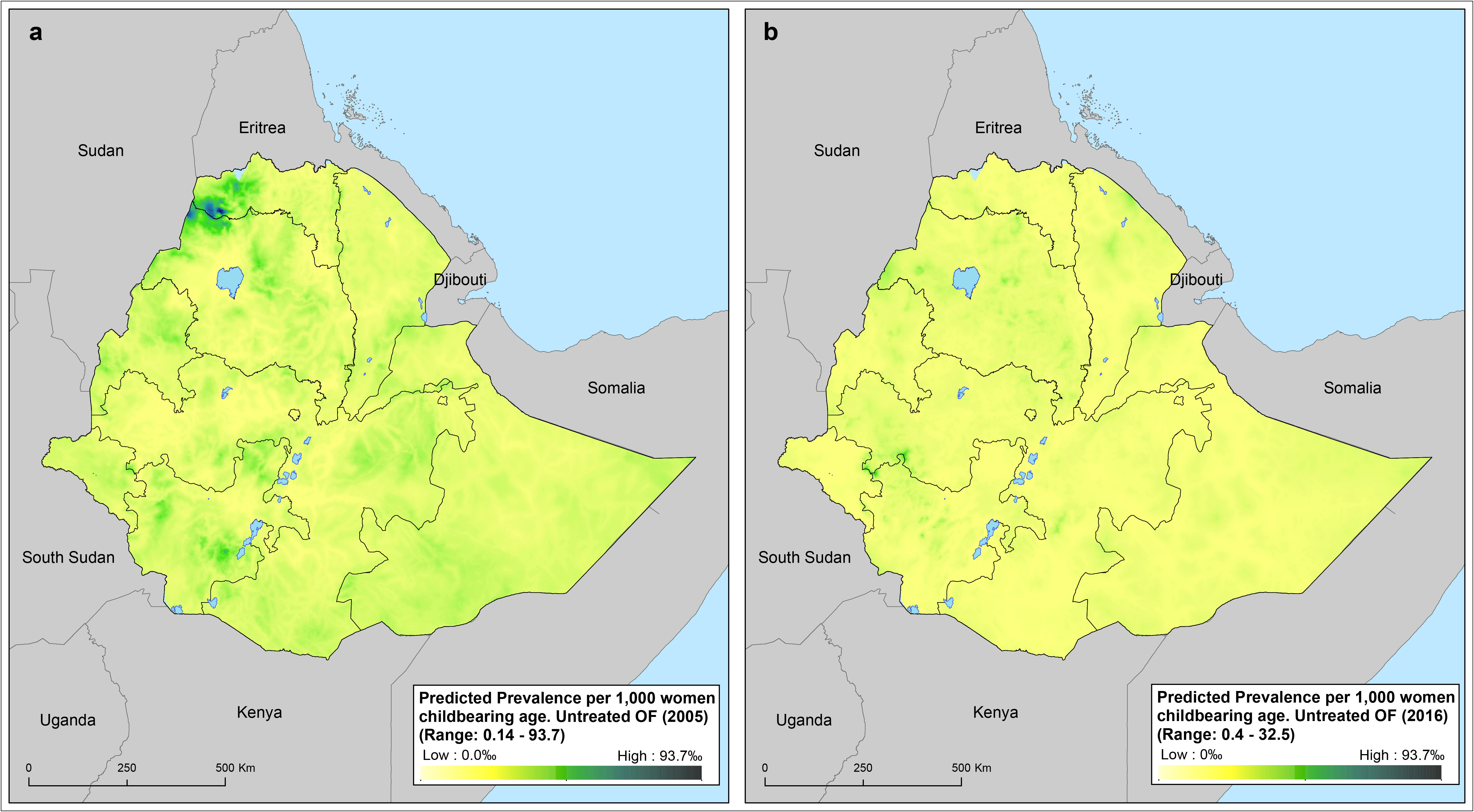
Predicted mean prevalence of untreated vaginal fistula in 2005 (A) and 2016 (B) across Ethiopia. Prevalence is provided in cases per 1,000 women of child bearing age.

**Figure 2.**
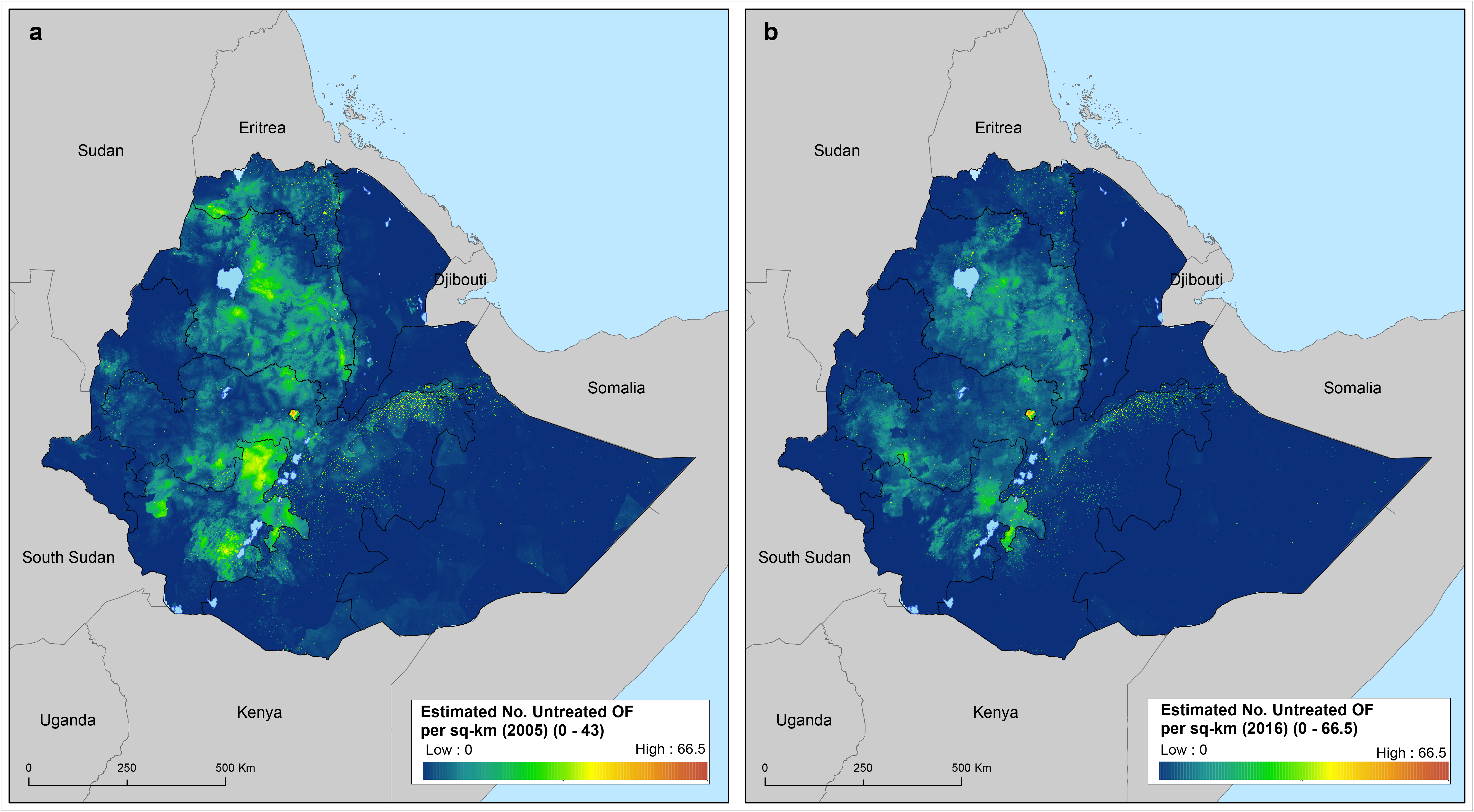
Estimated number of childbearing women (15-49 year-old) suffering from untreated vaginal fistula in 2005 (A) and in 2016 (B). Figures have been obtained using the estimated number of females aged between 15 and 49 per sq-km estimated by the WorldPop project.

In 2005 the prevalence of life time fistula in Ethiopia was 7.6 cases per 1000 WCA (95% CI: 3.7– 13.7). Prevalence varied between regions, from 2.7 (95% CI: 1.2–5.3) in Harari, to 15.4 (95% CI: 5.9– 32·4) in Tigray (Table 2). In 2016 the prevalence of lifetime fistula in Ethiopia was nearly halved the estimated prevalence in 2005, 3.4 cases per 1000 WCA (95% CI: 1.8–6.0), although the magnitude of reduction varied between regions. This was more significant in regions that showed the highest estimated prevalence such as Tigray (from 15.4 to 3.20 per 1000), Amhara (from 9.2 to 4.1 per 1000), Beneshangul Gumuz (from 9.4 to 3.8 per 1000) or Dire Dawa (from 9.1 to 2.9 per 1000). Harari, with the lowest estimated prevalence in 2005 (2.7, 95%CI: 1.2-5.3), however, showed no significant reduction in the estimated prevalence of lifetime fistula in 2016 (2.5, 95%CI: 1.3-4.4) (Table 2 & Table 3).

**Table 2.** Predicted mean prevalence and credible intervals of vaginal fistula in Ethiopia by 2005

|  | Predicted Prevalence per 1,000 |  |  |  |  |  |
| --- | --- | --- | --- | --- | --- | --- |
|  | Untreated vaginal Fistula |  |  | Lifetime vaginal Fistula |  |  |
|  | Lower Bound | Mean | Upper Bound | Lower Bound | Mean | Upper Bound |
| *Addis Ababa* | 0.46 | 1.66 | 4.34 | 1.65 | 4.93 | 9.76 |
| *Afar* | 1.96 | 3.73 | 6.73 | 2.70 | 4.64 | 7.47 |
| *Amhara* | 2.19 | 5.42 | 12.00 | 4.38 | 9.21 | 16.99 |
| *Beneshangul Gumuz* | 2.55 | 5.52 | 10.58 | 5.05 | 9.43 | 15.82 |
| *Dire Dawa* | 2.95 | 6.31 | 12.01 | 4.86 | 9.06 | 14.94 |
| *Gambella* | 2.16 | 4.84 | 10.60 | 3.95 | 7.78 | 15.20 |
| *Harari* | 0.65 | 1.84 | 4.35 | 1.20 | 2.70 | 5.27 |
| *Oromia* | 2.09 | 3.99 | 7.21 | 3.27 | 5.66 | 9.09 |
| *SNNP* | 2.59 | 5.84 | 11.94 | 4.53 | 8.86 | 15.79 |
| *Somali* | 2.36 | 4.50 | 8.43 | 3.09 | 5.39 | 8.57 |
| *Tigray* | 2.72 | 9.00 | 23.33 | 5.95 | 15.39 | 32.39 |
| **Total Averaged** | **2.06** | **4.79** | **10.14** | **3.69** | **7.55** | **13.75** |

**Table 3.** Predicted mean prevalence and credible intervals of vaginal fistula in Ethiopia by 2016

|  | Predicted Prevalence per 1,000 |  |  |  |  |  |
| --- | --- | --- | --- | --- | --- | --- |
|  | Untreated vaginal Fistula |  |  | Lifetime vaginal Fistula |  |  |
|  | Lower Bound | Mean | Upper Bound | Lower Bound | Mean | Upper Bound |
| *Addis Ababa* | 0.15 | 1.43 | 5.25 | 0.80 | 3.09 | 7.49 |
| *Afar* | 0.62 | 1.59 | 3.32 | 2.24 | 3.96 | 6.45 |
| *Amhara* | 0.87 | 2.27 | 4.67 | 2.24 | 4.11 | 6.74 |
| *Beneshangul Gumuz* | 0.73 | 1.70 | 3.37 | 2.26 | 3.80 | 5.95 |
| *Dire Dawa* | 0.34 | 0.85 | 1.70 | 1.72 | 2.95 | 4.62 |
| *Gambella* | 0.36 | 1.06 | 2.73 | 1.62 | 3.24 | 6.01 |
| *Harari* | 0.19 | 0.61 | 1.45 | 1.30 | 2.52 | 4.38 |
| *Oromia* | 0.56 | 1.38 | 2.75 | 2.05 | 3.49 | 5.43 |
| *SNNP* | 0.57 | 1.54 | 3.31 | 1.99 | 3.59 | 5.94 |
| *Somali* | 0.48 | 1.34 | 3.13 | 1.92 | 3.71 | 6.57 |
| *Tigray* | 0.37 | 1.23 | 3.16 | 1.53 | 3.20 | 6.01 |
| **Total Averaged** | **0.48** | **1.36** | **3.17** | **1.79** | **3.42** | **5.96** |

The prevalence of untreated VF symptoms was estimated as 4.8 cases per 1000 WCA (95% CI: 2.1– 10.1) in 2005 (Table 2). There has been a significant reduction to 1.4 cases per 1000 (95% CI: 0.5– 3.2) in 2016 (Table 3). The highest prevalence of untreated vaginal fistula in 2016 was found in Amhara region with 2.3 cases per 1000 WCA (95% CI: 0.9–4.7), while Harari had registered the lowest prevalence of untreated fistula in Ethiopia with 0.6 cases per 1000 WCA (95% CI: 0.2–1.4) (Table 3).

We estimated the number of WCA who had ever had VF symptoms to be 98,098 (95% CI: 49,819– 170,737) in 2005 (Table 4) and 72,533 (95% CI: 38,235–124,103) in 2016 (Table 5). We have also estimated the number of WCA with current VF symptoms to be 59,114 (95% CI: 26,580–118,158) in 2005 (Table 4) and 31,961(95% CI: 11,596–70,309) in 2016 (Table 5). Oromia, Amhara and SNNP bear the highest burden of WCA who had ever had VF symptoms and women who presently had symptoms of VF. The lowest burden of current symptoms of VF was predicted in Harari, Gambella and Dire Dawa (Tables 4 & 5).

**Table 4.** Estimated childbearing age women affected by vaginal fistula by 2005

|  | Estimated 19-49-y-old women 2005 |  |  |  |  |  |
| --- | --- | --- | --- | --- | --- | --- |
|  | Untreated vaginal Fistula |  |  | Lifetime vaginal Fistula |  |  |
|  | Lower Bound | Mean | Upper Bound | Lower Bound | Mean | Upper Bound |
| **Addis Ababa** | 314 | 1,339 | 3,773 | 1,281 | 4,254 | 8,801 |
| **Afar** | 556 | 1,065 | 1,883 | 816 | 1,416 | 2,265 |
| **Amhara** | 7,392 | 16,167 | 31,761 | 14,314 | 27,458 | 46,901 |
| **Beneshangul Gumuz** | 426 | 923 | 1,757 | 847 | 1,584 | 2,650 |
| **Dire Dawa** | 131 | 427 | 1,085 | 381 | 1,111 | 2,202 |
| **Gambella** | 129 | 288 | 637 | 239 | 475 | 930 |
| **Harari** | 15 | 46 | 103 | 37 | 87 | 165 |
| **Oromia** | 8,213 | 16,616 | 30,688 | 14,445 | 26,042 | 42,552 |
| **SNNP** | 6,333 | 15,051 | 30,974 | 12,134 | 24,599 | 43,999 |
| **Somali** | 1,663 | 3,228 | 6,065 | 2,210 | 3,915 | 6,369 |
| **Tigray** | 1,408 | 3,965 | 9,432 | 3,114 | 7,157 | 13,902 |
| **Total** | **26,580** | **59,114** | **118,158** | **49,819** | **98,098** | **170,737** |

**Table 5.** Estimated childbearing age women affected by vaginal fistula by 2016

|  | Estimated 19-49-y-old women 2016 |  |  |  |  |  |
| --- | --- | --- | --- | --- | --- | --- |
|  | Untreated vaginal Fistula |  |  | Lifetime vaginal Fistula |  |  |
|  | Lower Bound | Mean | Upper Bound | Lower Bound | Mean | Upper Bound |
| **Addis Ababa** | 117 | 1,301 | 4,928 | 733 | 3,261 | 8,475 |
| **Afar** | 186 | 473 | 995 | 805 | 1,410 | 2,267 |
| **Amhara** | 4,250 | 11,200 | 23,415 | 11,490 | 21,308 | 35,112 |
| **Beneshangul Gumuz** | 134 | 307 | 595 | 455 | 759 | 1,175 |
| **Dire Dawa** | 27 | 234 | 826 | 189 | 705 | 1,777 |
| **Gambella** | 37 | 98 | 230 | 158 | 295 | 517 |
| **Harari** | 7 | 48 | 172 | 58 | 178 | 431 |
| **Oromia** | 3,882 | 9,822 | 19,975 | 13,521 | 23,674 | 37,784 |
| **SNNP** | 2,103 | 5,813 | 12,487 | 7,118 | 13,401 | 22,716 |
| **Somali** | 463 | 1,298 | 2,998 | 1,994 | 3,828 | 6,689 |
| **Tigray** | 390 | 1,366 | 3,688 | 1,715 | 3,715 | 7,159 |
| **Total** | **11,596** | **31,961** | **70,309** | **38,235** | **72,533** | **124,103** |

Based on our models, the estimated prevalence of untreated vaginal fistula only exceeded 20 per 1000 childbearing age women in two districts of both Tigray and Amhara regions in 2005, and no single district was estimated to exceed this prevalence in 2016 (Fig 3). When exploring the burden of vaginal fistula by district, we found that the number of districts that were estimated to have more than 200 cases of untreated vaginal fistula dropped from 54 in 2005 to 6 in 2016. The majority of the districts with untreated cases were located in Amhara, Tigray, SNNP and eastern Oromia Regions (Fig 4).

**Figure 3.**
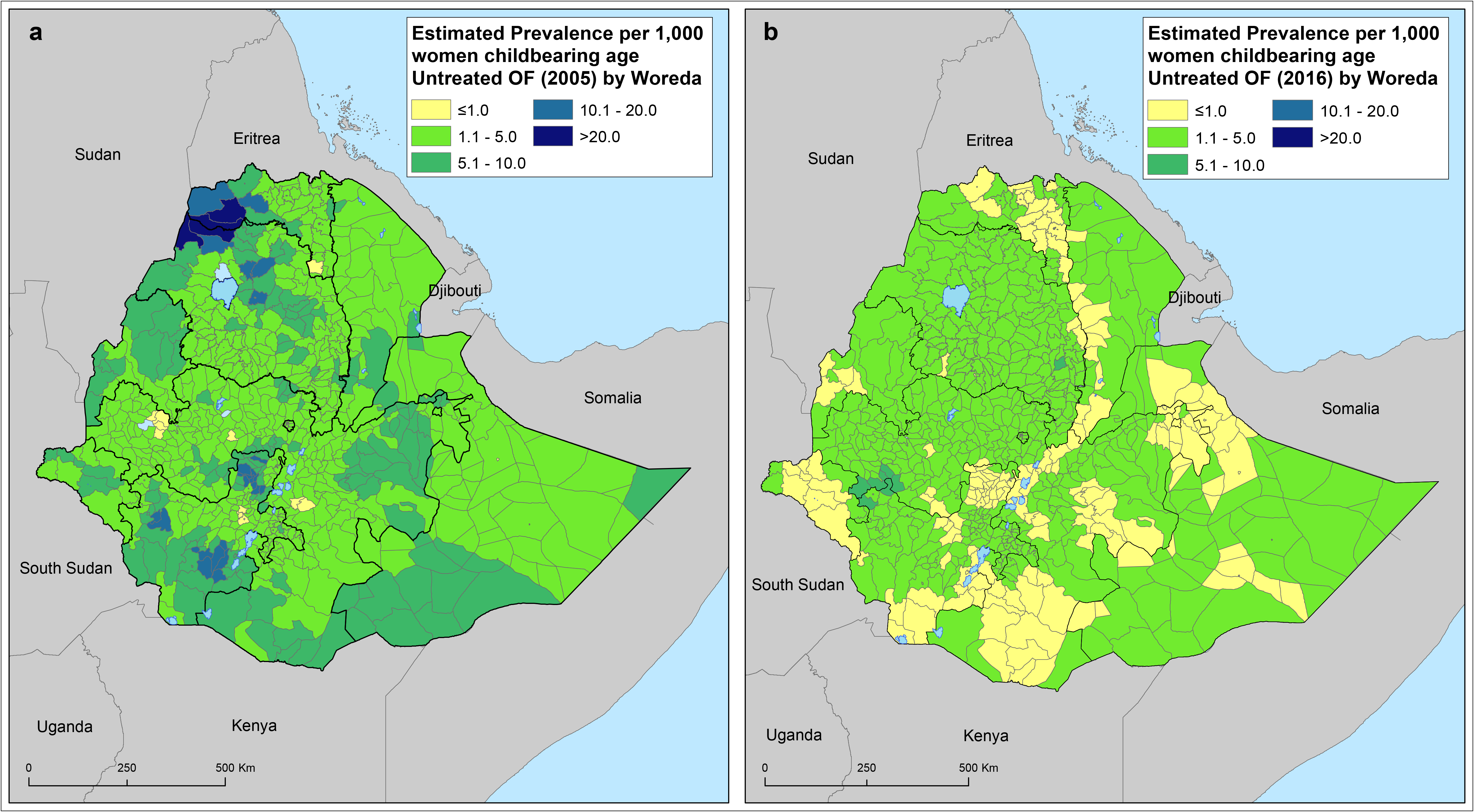
Predicted mean prevalence of untreated vaginal fistula by district in 2005 (A) and 2016 (B) across Ethiopia. Prevalence is provided in cases per 1,000 women of child bearing age.

**Figure 4.**
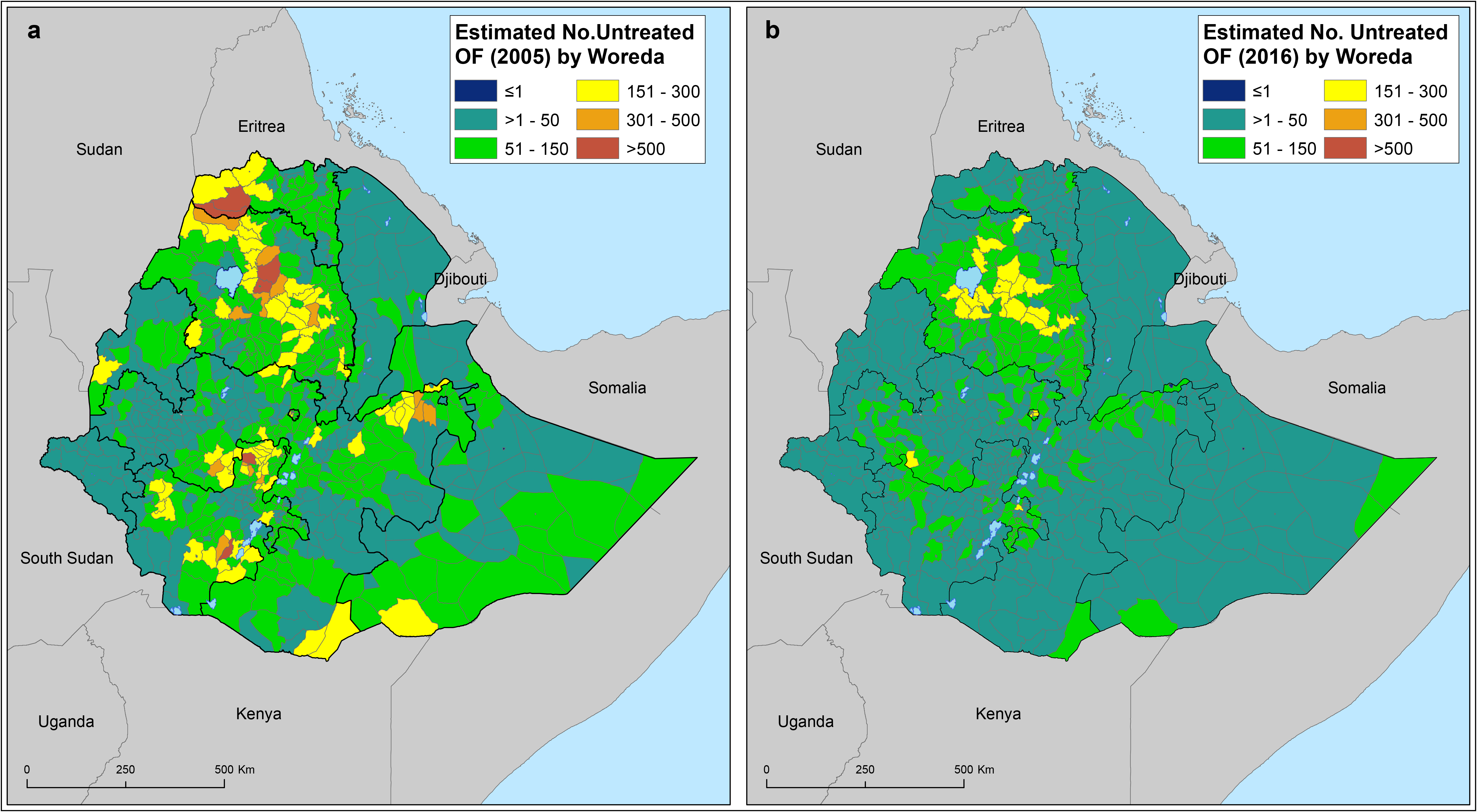
Estimated number of childbearing women (15-49 year-old) suffering from lifetime vaginal fistula by district in 2005 (A) and in 2016 (B). Figures have been obtained using the estimated number of females aged between 15 and 49 per sq-km estimated by the WorldPop project.

In 2016 lifetime VF prevalence of ranged from 2.3 per 1000 in Endegagn district (SNNP Region) to 6.7 per 1000 in Sigmo district (Oromia Region; Supplementary Information). In 2005 more than 10 per 1000 WCA experienced life time VF in Ethiopia in 105 districts in Ethiopia. In 2016 in none of the district registered > 10 per 1000 WCA lifetime VF (Supplementary Information). In 2005 more than 10 per 1000 women of reproductive age experienced life time fistula in Ethiopia in 29 districts. In 2016 in none of the districts > 10 per 1000 WCA life time VF was registered (Fig 3).

### Model validation

We provide in a supplementary information (Figures S9 & S10) the output of our validation procedure for the binomial mixed models as described under methods section. We simulated 500 data set from the fitted model and calculated the mean and standard deviation. We then compared the distribution of these two statistics with the ones observed in the data. For both lifetime and untreated OF models, there is agreement between the simulation from our models and the metrics observed in the empirical data, indicating a good predictive performance of the fitted models.

## Discussion

This is the first geostatistical analysis to estimate the burden of VF at district level in a developing country setting. We used large scale survey data in combination with socio-economic, demographic and health related continuous data to obtain these estimates. The point and life time prevalence of VF exhibited considerable heterogeneity across districts. The prevalence of symptoms of vaginal fistula although decreasing, still is considerably high. Districts with high prevalence are clustered in Amhara, Oromia and SNNP Regions. According to our estimates, in 2016, there were 31,961 (95% CI: 11,596–70,309) untreated cases of VF in Ethiopia, with majority of the cases in Oromia, Amhara and SNNP and with reduction from 2005 of 72,533 (95% CI: 38,235–124,103) untreated cases.

The point prevalence of VF, 1.4 per 1000 WCA, is higher than the prevalence reported by Ballard ^9^ and colleagues (0.2 untreated fistula prevalence per 1000 WCA), and consistent with the 1.5 per 1000 WCA reported by Muleta and colleagues ^10^ and global estimates for Sub-Saharan (1-1.6 per 1000 WCA) ^7,8^. The point prevalence of VF estimates in 2005 was slightly lower than a previous estimation which used the same data sets ^7^. The difference between the two estimations is attributed to the definition of treated cases. We assumed all women who reported, “having received treatment”, regardless of the outcome of the treatment, as treated cases. The previous analysis considered those women who did not report the outcome of the treatment as untreated cases.

We estimated that there were 31,961 (95% CI: 11,596–70,309) cases of untreated women with vagina fistula symptoms and 72,533 (95% CI: 38,235–124,103) lifetime vaginal fistula in 2016. There is a significant reduction from 2005 figures. The reduction in the prevalence of fistula between 2005 and 2016 could be explained by the introduction of health extension workers (HEW) at the community level, which facilitated identification, and referral of women with symptoms of vaginal fistula. It could also be attributed to the government’s ambitious National Action Plan launched in 2014 to eliminate fistula in six years’ time ^11^.

The results also clearly show that vagina fistula is a ubiquitous and spatially heterogeneous health problem in Ethiopia. There is clear clustering of vagina fistula symptoms in some districts. Districts with high burden of untreated vagina fistula symptoms include Fareta, Ebenat and Dessie Zuria at the Amhara region. The number of untreated VF cases varies between districts from 0 to 291. Arguably, districts in the central Amhara, Oromia, and northern SNNP are the areas, which bear the highest burden of VF. These districts required more attention to reduce the burden of VF in the country. Districts surrounding Hawassa have to some extent access to emergency obstetric care. Thus, it is important to investigate the reason why these districts have high prevalence despite having access to emergency obstetric care. To achieve the elimination of obstetric fistula, the highest burden districts should primarily be targeted to clear the existing backlog of VF symptoms. In addition, investigating district-specific determinants with focus to high burden districts is warranted.

The strength of our analysis include use of large surveys with well powered sample size, covering large geographical areas and population groups and the use of standardize questionnaire for measuring VF. We generated estimates at district level. District level data allows benchmarking of subnational administrative units, which will help identify best and worst performing districts and design tailored interventions for the different scenarios. However, we must acknowledge some limitations of our analysis. First, VF symptoms is a highly stigmatizing condition so that substantial number of women suffering from this condition might have been expelled from their homes due to ostracism by their families and seek refuge in long term care facilities dedicated to patients with fistula. Considering that DHS is a household based survey, the aforementioned scenarios may lead to many cases not being recorded and therefore the point prevalence be underestimated ^7,12^. Second, the DHS surveys only target WCA for self-reported vaginal fistula, although previous studies have documented low incidence of vaginal fistula among women older than age 50 years and those younger than 15 years ^7^. This can also lead to underestimating the prevalence of VF. Finally, measurement bias could be another limitation ^13,14^ nonetheless, we have accounted for sensitivity and specificity of the DHS questionnaire based on previous studies ^7,13,15^. However, we do not believe that these limitations diminish the overall interpretations, program and policy implications and actions based on our findings.

Our findings also highlight the importance of tracking symptoms of vaginal fistula at granular level for monitoring and programme evaluation. These can be achieved through collection such data through HEW at regular interval or through routine community based surveillance at district level. Our findings have several importance for policy and programming. First, the findings here will be of help in benchmarking district level burden of VF, support monitoring current and future interventions against this condition. Second, identifying districts with high burden of VF will help prioritize target areas for intervention and optimize resource allocation. By addressing district with high burden districts, the country can speed up the elimination of VF. In addition, program planners could identify district level interventions based of the burden of VF.

Third, identification of district with low and high burden VF would open an important research avenue for social and behavioural scientist. Comparative studies on identifying contextual and sociocultural factors that contribute to low and high prevalence of VF will have important implications in addressing the root causes of VF as it has been in other studies ^16–18^. Finally, we have demonstrated how spatial analysis can be used to estimate geographical disparity of the burden of vaginal fistula across Ethiopia. Similarly, this framework has the potential to be effectively integrated in the national health information system to track burden of other health conditions in the country.

In conclusion, our estimates have proven a sharp reduction of the VF between 2005 and 2016 in Ethiopia ^7^. The number of districts with >200 cases of untreated vaginal fistula reduced from 54 in 2005 to 6 in 2016. The difference could be due to the improved health system and maternal health services, the increased availability and accessibility of surgical services and specific efforts to end fistula in the country. Despite the progress made, there are still significant number of untreated cases in the country. Therefore, decentralized efforts targeting high burden districts is required to stop fistula being a public health problem. Elimination of obstetric fistula by 2020 as stipulated in the Health Sector Transformation Plan of Ethiopia ^11^ is off track and effort should be intensified on prevention interventions and treatment of the existing ones. Strong surveillance system to identify old and new cases of vaginal fistula and linking them to the needed surgical services should be strengthened by focusing on districts with high burden. There is a favourable momentum in Ethiopia to eliminate fistula, identification and referral of women with vaginal fistula is tremendously facilitated by the existence of HEW ^19,20^. The huge boost in the number of health centres, trained midwives help manage deliveries and avoid obstructed labour that causes obstetric fistula; expansion of hospitals and trained surgical officers over the years have expanded access to fistula surgery ^20,21^. Social integration of those who suffered fistula that addresses the physical, mental, social, legal and psychological needs of women with this condition is also of paramount importance.

## Methods

### Data sources

The 2005 and 2016 Ethiopian DHS (EDHS) ^6,22^ provide the largest, most comprehensive vaginal fistula assessment in Ethiopia. The surveys are representative at national and regional level in Ethiopia ^6,22^. The surveys included questions that were intended to provide estimates on vaginal fistula ^6,22^.

Administratively, Ethiopia is divided into nine geographical regions and two administrative cities ^6,22^. The sample for the EDHS was designed to provide estimates of key indicators for the country, for urban and rural areas separately, and for each of the nine regions and the two administrative cities 6,22. Each region was stratified into urban and rural areas, yielding 21 sampling strata. Samples of enumeration areas (EAs) were selected independently in each stratum in two stages^6,22^. Implicit stratification and proportional allocation were achieved at each of the lower administrative levels by sorting the sampling frame within each sampling stratum before sample selection, according to administrative units in different levels, and by using a probability proportional to size selection at the first stage of sampling^6,22^.

In the 2005 EDHS, a representative sample of approximately 14,500 households from 540 clusters were selected^22^. The sample was selected in two stages. In the first stage, 540 clusters (145 urban and 395 rural) (Supplementary Information, Fig S1 & S2) were selected from the list of EAs from the 1994 Population and Housing Census (PHC) sample frame^22^. The 2016 DHS was conducted from January 18, 2016, to June 27, 2016 in a total of 645 EAs (202 in urban areas and 443 in rural areas) (Supplementary Information; Fig S3 & S4)^6^. They were selected with probability proportional to EA size (based on the 2007 PHC) and with independent selection in each sampling stratum ^6^. All women age 15-49 who were either permanent residents of the selected households or visitors who stayed in the household the night before the survey were eligible to be interviewed ^6^.

### Measurement of outcomes

We aimed to estimate the district-level lifetime and untreated vaginal fistula among WCA and to explore changes overtime. The 2005 and 2016 EDHS included a specific thorough questionnaire of 11 questions intended to diagnose vaginal fistula and to ascertain the severity of the condition ^6,22^. Women were asked if they had experienced vaginal fistula as defined by *‘constant leakage of urine or stool from your vagina during the day and night?’* ^6,22^ Those who reported suffering from vaginal fistula were asked if they had ever been treated for this condition ^6,22^. We assessed two main estimates of prevalence following the approach used by Maheu-Giroux and colleagues ^7^: i) lifetime prevalence of fistula, as the proportion of respondents who reported having ever had symptoms of vaginal fistula, and ii), point prevalence (untreated fistula) of fistula symptoms by excluding those women who sought treatment for vaginal fistula ^7^.

## Data Analysis

### Sources of covariates

As it is detailed below, models for lifetime and untreated vaginal fistula were constructed using a set of covariates that may potentially be associated with the risk for vaginal fistula ^7,15,23^. We considered covariates that were available as continuous gridded maps for the whole country. From the Spatial Data DHS Program ^24^, we obtained modelled surfaces of the following covariates indicated to be associated with a higher risk for vaginal fistula: i) percentage of literate women ^23,25^, ii) proportion of live births in the five (or three) years preceding the survey delivered at a health facility ^4,23,26^, iii) proportion of women using any modern method of contraception and iv) proportion of women who had a live birth in the five (or three) years preceding the survey who had four or more antenatal care visits 15,23 (Figures 5S & 6S, Supplementary information). These modelled surfaces are available for Ethiopia at a spatial resolution of 5km × 5km and have been produced using standardized geostatistical methods, publicly available EDHS data collected in 2016, and a standardized set of covariates ^27^.

To account for disparities between rural and urban, we used gridded maps of urban accessibility obtained from the European Commission Joint Research Centre Global Environment Monitoring Unit (JRC) ^28^ and from the Malaria Atlas project ^29,30^ for 2000 and 2015, respectively (Figure S6, Supplementary information). Accessibility is time (minutes) taken to travel from a grid cell in the map to a city using land-based travel and employing the minimum cost. Cost-distance or impedance was determined by a friction surface that, although generated using different sources of data for the 2000 and 2015 accessibility models, in both cases contained information on the transport network and environmental and political factors that affect travel times between locations. Further details on the generation of this friction surfaces are provided elsewhere ^28,30^. We later used the friction surface from 2015 accessibility model to construct a gridded map of distances (kilometres) to the nearest health facility (Figure S5, Supplementary information). For this, we used a list of health facilities with their geographic coordinates downloaded from the Humanitarian Data Exchange database ^31^. This dataset, provided by the Ethiopian Ministry of Health, was created in 2012 and is regularly updated. The cost-distance tool available at the Spatial Analyst tool from ArcGIS 10.5 (ESRI Inc., Redlands CA, USA) was used to estimate the nearest distance from every cell grid in a 5km × 5km resolution gridded map from Ethiopia to the nearest health facility.

Population density estimates were obtained from the WorldPop project ^32^ for 2005 and 2015. These estimates were used to classify areas as urban (population densities ≥1,000 persons/km^2^), peri-urban (>250 persons/km^2^ within a 15 km distance from urban extents edge)or rural areas (250 persons/km^2^ and/or >15 km from the urban extents edge), ^2,33^ (Figure S7, Supplementary information).

Finally, night-light emissivity for 2005 and 2013 (most recent available) captured by the Operational Linescan System instrument on board a satellite of the Defence Meteorological Satellite Programme was used as a proxy measure of poverty across Ethiopia ^34,35^ (Figure S7, Supplementary information).

This instrument measures visible and infrared radiation emitted at night-time, resulting in remote imagery of lights on the ground. This information has been correlated with gross domestic product in developed countries ^35–37^ and, although far from precise, would provide an indirect measure of poverty in developing countries ^38,39^.

Input grids were resampled to a common spatial resolution of 1 square-km using a bilinear interpolation approach and clipped to match the geographic extent of a map of Ethiopia, and eventually aligned to it^35^. Raster manipulation and processing was undertaken using *raster* package in in the R software environment ^35,39,40^ and final map layouts created with ArcGIS 10.5 software (ESRI Inc., Redlands CA, USA).

We followed a model selection procedure to identify an optimal suite of covariates to include in the fixed effects part of the binomial models. In order to reduce any potential collinearity and confounding effects, we first grouped the variables and use a formal model selection criterion to select one variable within each of the groups^41^ (Tables S1 & S2, Supplementary information). Variables showing some level of correlation or have similar nature (i.e. accessibility and urbanization-related variables) were grouped together. Within each group, we investigated the relationship between the risks for untreated and lifetime vaginal fistula and each potential explanatory variable by fitting univariate models relating the logit of the outcomes to each of the variables ^41^(Tables S3 & S4, Supplementary information). We compared the univariate models in terms of the Akaike Information Criterion (AIC) and selected the model with the lowest value of AIC within each group of variables (equation 1). AIC is defined as

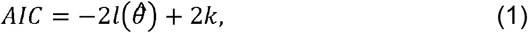

where 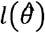 is the maximum log-likelihood function and *k* is the number of parameters.

### Modelling analysis

We estimated district-level prevalence of lifetime and untreated vaginal fistula for 2005 and 2016 across Ethiopia using a binomial mixed model, accounting for fixed effects (covariates) and random effects, for potential unexplained variability between regions^42^. We chose this method over other analytic approaches based on the absence of spatial structure on the prevalence of lifetime and untreated fistula prevalence, explored by fitting an empirical variogram ^42^ (Figure S8, Supplementary information). Our model used the prevalence estimates and covariates for smoothing the prediction (equation 2). Briefly, let *Y_ki_* denote the random variable associated with the number of positively detected cases of lifetime and untreated vaginal fistula at community location *x_ki_* with *k* indicating the region^42^. We then modelled *Y_ki_* using a binomial mixed model with probability of having lifetime fistula *p(x_ki_)* such that^35^

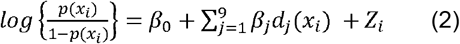

where the *d*_*j*_(*x*_*i*_) are geo-referenced covariates (described previously) and the *Z_k_* are independent and identically distributed zero-mean Gaussian variables with variance *σ*^2^. We fitted the model using the *lme4* package in the R software environment ^42,43^. Prior to construct the models, observed lifetime and untreated prevalence of vaginal fistula at location level were adjusted by assuming a sensitivity and specificity of fistula questionnaires common to all surveys of 97% and 99% respectively, as suggested by some authors ^7,15^. Adjustment was implemented as suggested by Diggle (2011) ^44^.

Models misspecification and goodness-of-fit were tested by simulating dataset (n= 500) from the fitted models, and eventually comparing summary statistics computed on the simulated data with the empirical ones. As result of good model performance, we expect that our model is able to replicate the characteristics of the observed prevalence.

Fitted models were applied on the selected covariates to produce continuous predictions of lifetime and untreated vaginal fistula prevalence, predicted mean prevalence and corresponding 95% credible intervals, at 1km^2^ spatial resolutions. Gridded maps of both population density and age structure were obtained from the WorldPop project ^42,45,46^. We used this gridded population surface for computing the estimates of affected population by pixel by multiplying prevalence of lifetime and untreated vaginal fistula in each square-km area with the corresponding population of women of childbearing age (15-49 year-old women) at the same spatial resolution^42^. Using the surfaces, number of women with lifetime and untreated vaginal fistula by district in 2005 and 2016 were extracted.

The National Ethics Review Committee of the Ethiopia Science and Technology Commission in Addis Ababa, Ethiopia and the ORC Macro Institutional Review Board in Calverton, USA approved the DHS 2005 study protocol^22^. The Federal Democratic Republic of Ethiopia Ministry of Science and Technology and the Institutional Review Board of ICF International reviewed and approved the DHS Ethiopia 2016 survey protocol^6^. Informed consent was obtained from all participants and/or their legal guardians. All experiments involving human subjects and samples in this study were performed in accordance with relevant guidelines and regulations^6,22^.

### Data availability

The data used in this publication are publicly available from https://dhsprogram.com/data/.

## Acknowledgements

We thank the permission granted by MEASURE DHS to use the DHS datasets.

KD is supported by the Wellcome Trust to as part of his International Intermediate Fellowship [grant number 201900].

## Authors’ contributions

KD and JC conceptualized the study. KD, CF, TD and JC analysed the data. KD Wrote the first draft of the manuscript. CF, TD, SB, AD, MA and JC Contributed to the writing of the manuscript. All agree with the manuscript’s results and conclusions. All authors have read, and confirm that they meet, ICMJE criteria for authorship.

## References

1 Biadgilign S., Lakew Y., Reda A.A. & Deribe K. A population based survey in Ethiopia using questionnaire as proxy to estimate obstetric fi stula prevalence: results from demographic and health survey Reprod Health 10, doi:10.1186/1742-4755-10-14. (2013).

2 Arrowsmith S., Hamlin E.C. & Wall L.L. Obstructed labor injury complex: obstetric fistula formation and the multifaceted morbidity of maternal birth trauma in the developing world. Obstet Gynecol Surv 51, 568–574 (2013).

3 Wall L.L. Obstetric vesicovaginal fistula as an international public-health problem. Lancet 368, 1201–1209 (2013).

4 Wall L.L., Arrowsmith S.D., Briggs N.D., Browning A. & Lassey A. The obstetric vesicovaginal fistula in the developing world. Obstet Gynecol Survey 60, S3–51 (2013).

5 UNFPA. & Campaign to End Fistula. http://endfistula.org/campaign (Accessed: 01 February 2019).

6 Central Statistical Agency (CSA) [Ethiopia]. & ICF. Ethiopia Demographic and Health Survey 2016. (Addis Ababa, Ethiopia, and Rockville, Maryland, USA, 2016).

7 Maheu-Giroux M. et al. Prevalence of symptoms of vaginal fistula in 19 sub-Saharan Africa countries: a meta-analysis of national household survey data. Lancet Glob Health 3, e271–278 (2013).

8 Adler A.J., Ronsmans C., Calvert C. & Filippi V. Estimating the prevalence of obstetric fistula: a systematic review and meta-analysis. BMC Pregnancy Childbirth 13, 246, doi:10.1186/-1471-2393-13-246. (2013).

9 Ballard K., Ayenachew F., Wright J. & Atnafu H. Prevalence of obstetric fistula and symptomatic pelvic organ prolapse in rural Ethiopia. Int Urogynecol J 27, 1063–1067 (2013).

10 Muleta M., Fantahun M., Tafesse B., Hamlin E.C. & Kennedy R.C. Obstetric fistula in rural Ethiopia. East Afr Med J 84, 525–533 (2013).

11 Federal Democratic Republic of Ethiopia Ministry of Health. Health Sector Transformation Plan (HSTP). Addis Ababa, Ethiopia. (2013).

12 Muleta M, Hamlin EC, Fantahun M, Kennedy RC & Tafesse B. Health and social problems encountered by treated and untreated obstetric fistula patients in rural Ethiopia. J Obstet Gynaecol Can 30, 44–50 (2013).

13 Tunçalp Ö., Isah A., Landry E. & Stanton C.K. Community-based screening for obstetric fistula in Nigeria: a novel approach. BMC Pregnancy Childbirth 14, doi:10.1186/1471-2393-14-44. (2013).

14 Tunçalp Ö, Tripathi V, Landry E, Stanton CK & Ahmed S. Measuring the incidence and prevalence of obstetric fistula: approaches, needs and recommendations. Bull World Health Organ 93, 60–62 (2013).

15 Maheu-Giroux, M. et al. Risk factors for vaginal fistula symptoms in Sub-Saharan Africa: a pooled analysis of national household survey data. BMC Pregnancy and Childbirth 16, doi:10.1186/s12884-016-0871-6. (2013).

16 Fèvre E.M. et al. The origins of a new Trypanosoma brucei rhodesiense sleeping sickness outbreak in eastern Uganda. Lancet 358, 625–628 (2013).

17 Fischer C.S., Stockmayer G., Stiles J. & Hout M. Distinguishing the geographic levels and social dimensions of U.S. metropolitan segregation, 1960-2000. Demography 41, 37–59 (2013).

18 Sullivan A.B. et al. Are neighborhood sociocultural factors influencing the spatial pattern of gonorrhea in North Carolina? Ann Epidemiol 21, 245–252 (2013).

19 Hamlin C. & Fleck F. Giving hope to rural women with obstetric fistula in Ethiopia. Bull World Health Organ 91, 724–725 (2013).

20 Duby F. & Hailey J. AusAID Health Resource Facility Managed by HLSP in association with IDSS. Joint AusAID and USAID Review of Support to Hamlin Fistula Ethiopia (Ethiopia) Final report. https://reliefweb.int/sites/reliefweb.int/files/resources/joint-aus-us-gov-review-hamlin-fistula-ethiopia-final-report.pdf. (Accessed: 22 July 2019). (2013).

21 Balabanova D. et al. Good Health at Low Cost 25 years on: lessons for the future of health systems strengthening. Lancet 381, 2118–2133 (2013).

22 Central Statistical Agency [Ethiopia]. & ORC Macro. Ethiopia Demographic and Health Survey 2005. (Addis Ababa, Ethiopia and Calverton, Maryland, USA, 2006).

23 Andargie, A. A. & Debu, A. Determinants of obstetric fistula in Ethiopia. Afr Health Sci 17, 671–680 (2013).

24 Burgert-Brucker C.R., Dontamsetti T. & Gething P.W. The DHS Program’s Modeled Surfaces Spatial Datasets. Stud Fam Plann 49, 87–92 (2013).

25 Tebeu P.M., de Bernis L., Doh A.S., Rochat C.H. & Delvaux T. Risk factors for obstetric fistula in the Far North Province of Cameroon. Int J Gynaecol Obstet 107, 12–15 (2013).

26 Tebeu P.M. et al. Risk factors for obstetric fistula: a clinical review. Int Urogynecol J 23, 387–394 (2013).

27 Burgert-Brucker, C. R., Dontamsetti, T., Marshall, A. M. J. & Gething, P. W. Guidance for use of the DHS program modelled map surfaces: DHS Spatial Analysis Reports 14. (United States Agency for International Development (USAID), 2016).

28 Joint Research Centre. The European Commission’s science and knowledge service. Travel time to major cities: A global map of Accessibility. http://forobs.jrc.ec.europa.eu/products/gam/. (Accessed: 8 May 2018).

29 MAP. Malaria Atlas project. https://map.ox.ac.uk/ (Accessed: 8 May 2018).

30 Weiss, D. J. et al. A global map of travel time to cities to assess inequalities in accessibility in 2015. Nature 553, 333–336 (2013).

31 HDX. The Humanitarian Data Exchange. https://data.humdata.org/ (Accessed: 7 May 2018).

32 WorldPop. The WorldPop demography project. http://www.worldpop.org.uk/ (Accessed: 11 May 2018).

33 Pullan, R. L. & Brooker, S. J. The global limits and population at risk of soil-transmitted helminth infections in 2010. Parasit Vectors 5, 81, doi:10.1186/1756-3305-5-81 (2013).

34 Elvidge, C. D., Baugh, K. E., Kihn, E. A., Kroehl, H. W. & Davis, E. R. Mapping city lights with nighttime data from the DMSP operational linescan system. Photogrammetric Engineering and Remote Sensing 63, 727–734 (2013).

35 Deribe K. et al. Estimating the number of cases of podoconiosis in Ethiopia using geostatistical methods Wellcome Open Res 2, doi:10.12688/wellcomeopenres.12483.2 (2013).

36 Doll, C. N. H., Muller, J. P. & Morley, J. G. Mapping regional economic activity from night-time light satellite imagery. Ecological Economics 57, 75–92 (2013).

37 Ebener, S., Murray, C., Tandon, A. & Elvidge, C. C. From wealth to health: modelling the distribution of income per capita at the sub-national level using night-time light imagery. Int J Health Geogr 4, 5, doi:10.1186/1476-072X-4-5 (2013).

38 Noor, A. M., Alegana, V. A., Gething, P. W., Tatem, A. J. & Snow, R. W. Using remotely sensed night-time light as a proxy for poverty in Africa. Popul Health Metr 6, 5, doi:10.1186/1478-7954-6-5 (2013).

39 Deribe K. et al. Predicted distribution and burden of podoconiosis in Cameroon. BMJ Glob Health 3, e000730, doi:10.1136/bmjgh-2018-000730 (2013).

40 Bates M., Mächler M., Bolker B. & Walker S. Fitting linear mixed-effects models using lme4. Journal of Statistical Software 67, 1–48 (2013).

41 Moraga P. et al. Modelling the distribution and transmission intensity of lymphatic filariasis in sub-Saharan Africa prior to scaling up interventions: integrated use of geostatistical and mathematical modelling. Parasit Vectors 8, 560, doi:10.1186/s13071-015-1166-x. (2013).

42 Deribe K. et al. Geographical distribution and prevalence of podoconiosis in Rwanda: a cross-sectional country-wide survey. Lancet Glob Health 7, e671–e680.

43 Bates M, Mächler M, Bolker B & Walker S. Fitting linear mixed-effects models Using lme4. Journal of Statistical Software 67, 1–48 (2013).

44 Diggle, P. J. Estimating Prevalence Using an Imperfect Test. Epidemiology Research International 2011, doi:10.1155/2011/608719 (2013).

45 Tatem A.J., Noor A.M., von Hagen C., Di Gregorio A. & Hay S.I. High resolution settlement and population maps for low income nations: combining land cover and national census in East Africa. PLoS One 2, e1298, doi:10.1371/journal.pone.0001298 (2013).

46 Linard C. et al. Population distribution, settlement patterns and accessibility across Africa in 2010. PLoS ONE 7, e31743, doi:10.1371/journal.pone.0031743 (2013).

