## Supplementary Material 1 for "Measuring the spatial heterogeneity on the reduction of vaginal fistula burden in Ethiopia between 2005 and 2016"

**Figure S1** Distribution of selected clusters for 2005 Demographic and Health Surveys and prevalence of obstetric fistula across Ethiopia.

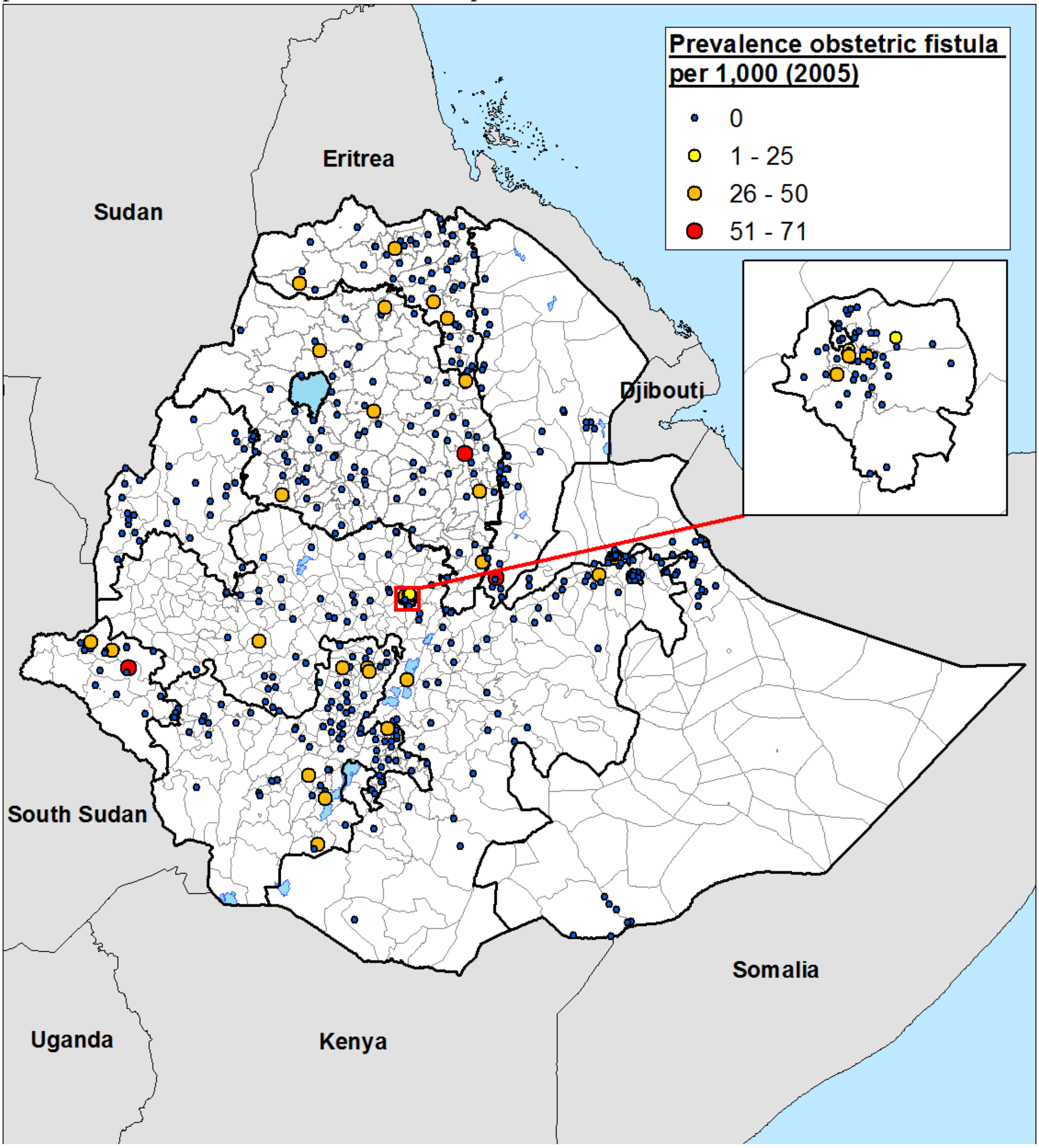

**Figure S2** Distribution of selected clusters for 2005 Demographic and Health Surveys and prevalence of lifetime obstetric fistula across Ethiopia.

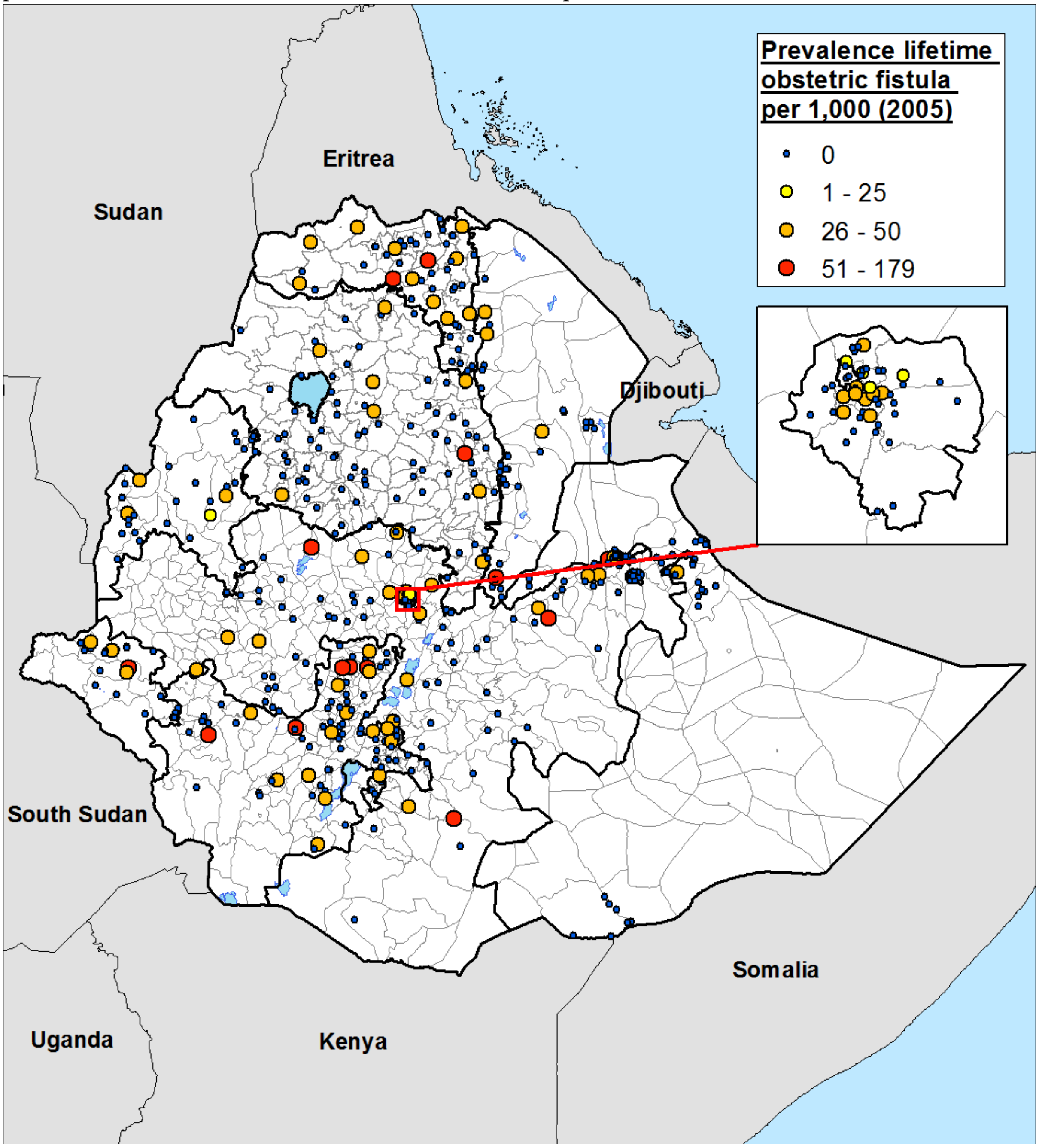

**Figure S3** Distribution of selected clusters for 2016 Demographic and Health Surveys and prevalence of obstetric fistula across Ethiopia.

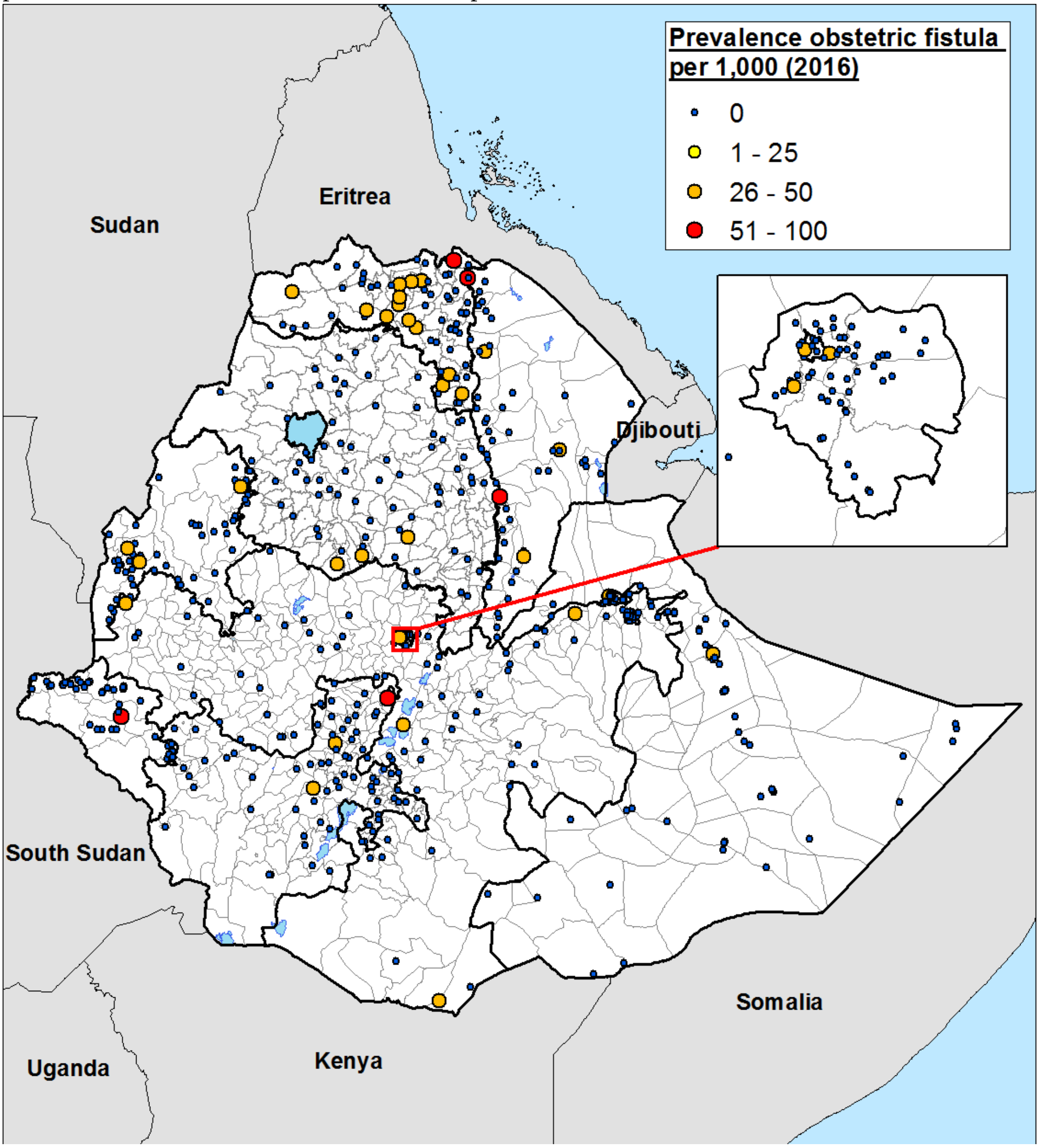

**Figure S4** Distribution of selected clusters for 2016 Demographic and Health Surveys and prevalence of lifetime obstetric fistula across Ethiopia.

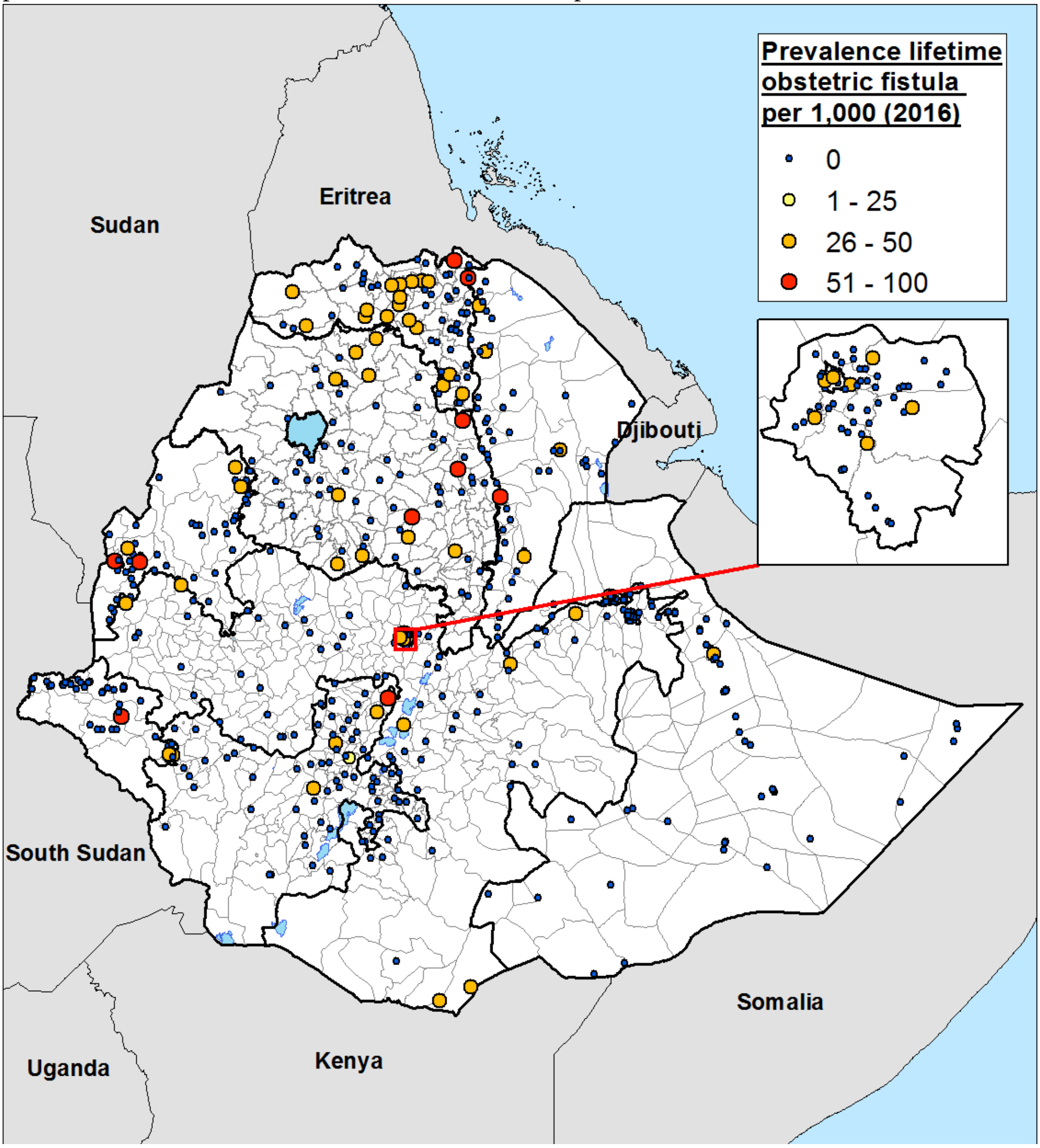

**Figure S5** Variables used to model prevalence of untreated and lifetime obstetric fistula, and to calculate estimates of childbearing-age population potentially affected in Ethiopia in 2005 and 2016. A) Cost distance nearest health facility (HF), B) Friction surface used to estimate nearest distance to HF, C) Proportion women using modern contraception methods, and D) Proportion literate women.

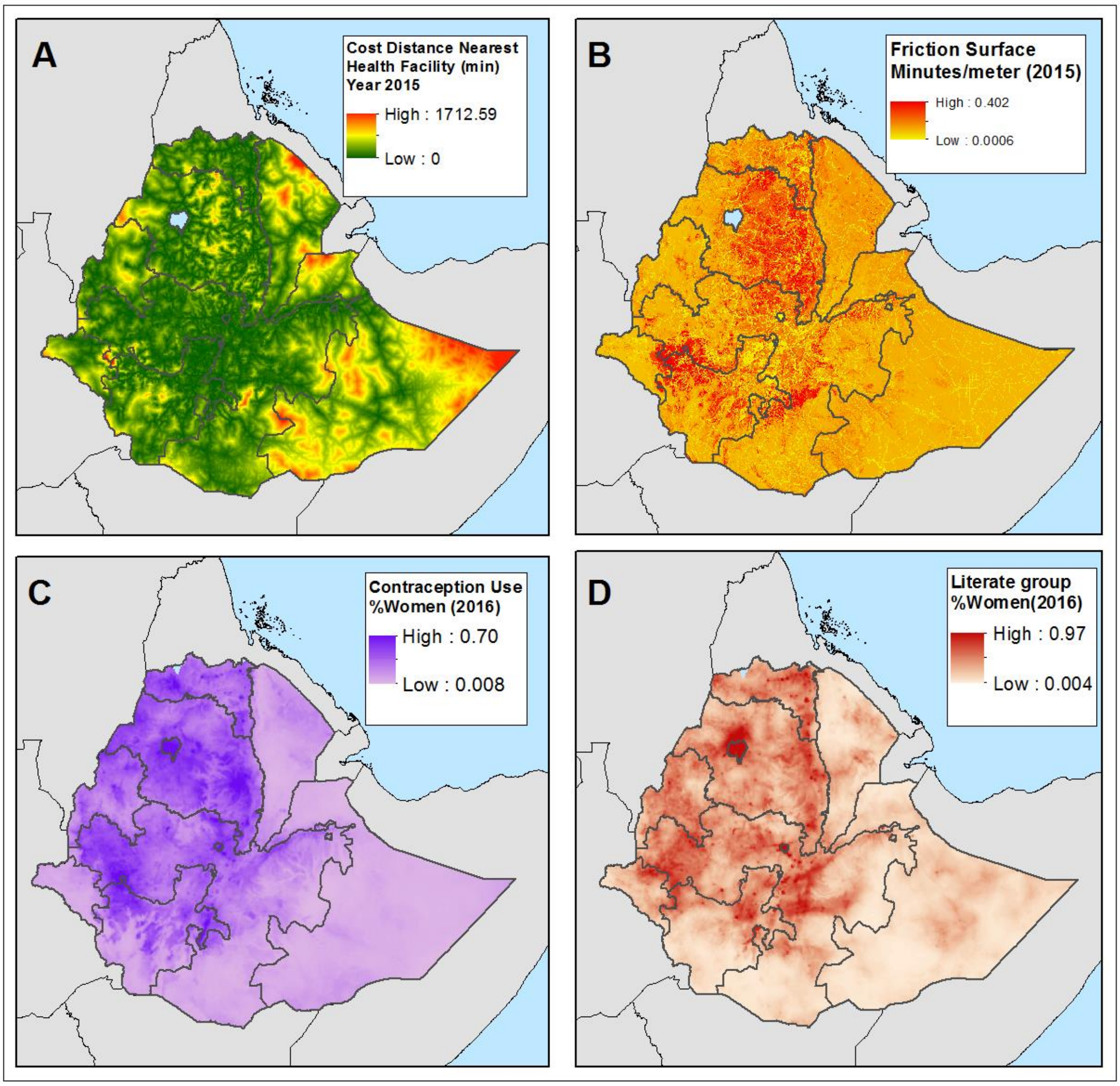

**Figure S6** Variables used to model prevalence of untreated and lifetime obstetric fistula, and to calculate estimates of childbearing-age population potentially affected in Ethiopia in 2005 and 2016. A) Accessibility (min) to nearest town (2000), B) Accessibility (min) to nearest town (2015), C) Proportion women having completed four antenatal visits, D) Proportion women delivering at a health facility (HF), E) Estimated childbearing-age population (19-49 year-old) in 2005, and F) Estimated childbearing-age population (19-49 year-old) in 2015.

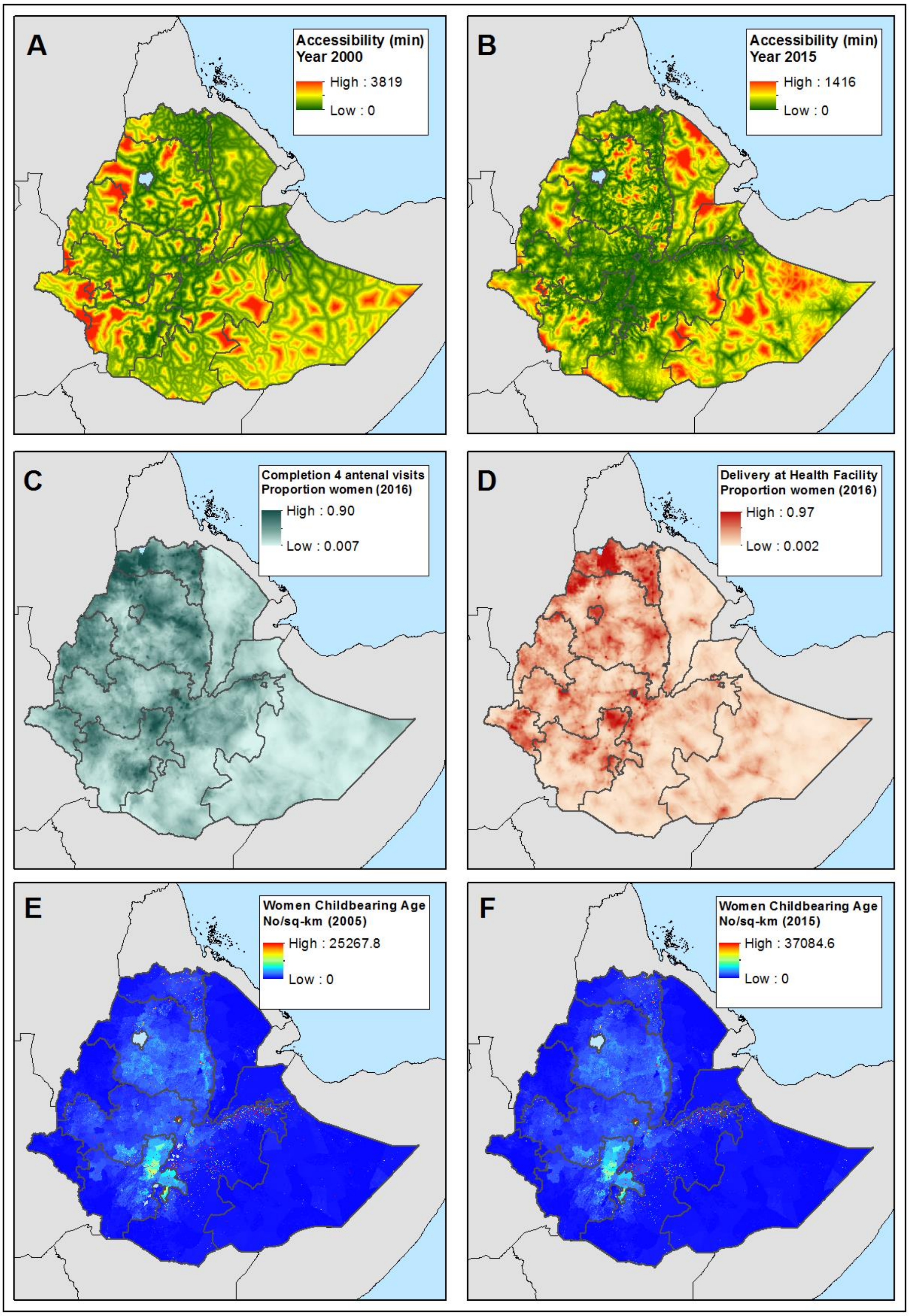

**Figure S7** Variables used to model prevalence of untreated and lifetime obstetric fistula, and to calculate estimates of childbearing-age population potentially affected in Ethiopia in 2005 and 2016. A) Night-light emissivity (2005), B) Night-light emissivity (2015), C) Distance (km) stable night-lights in 2005, D) Distance (km) stable night-lights in 2013, E) Level of Urbanization in 2005, and F) Level of Urbanization in 2015.

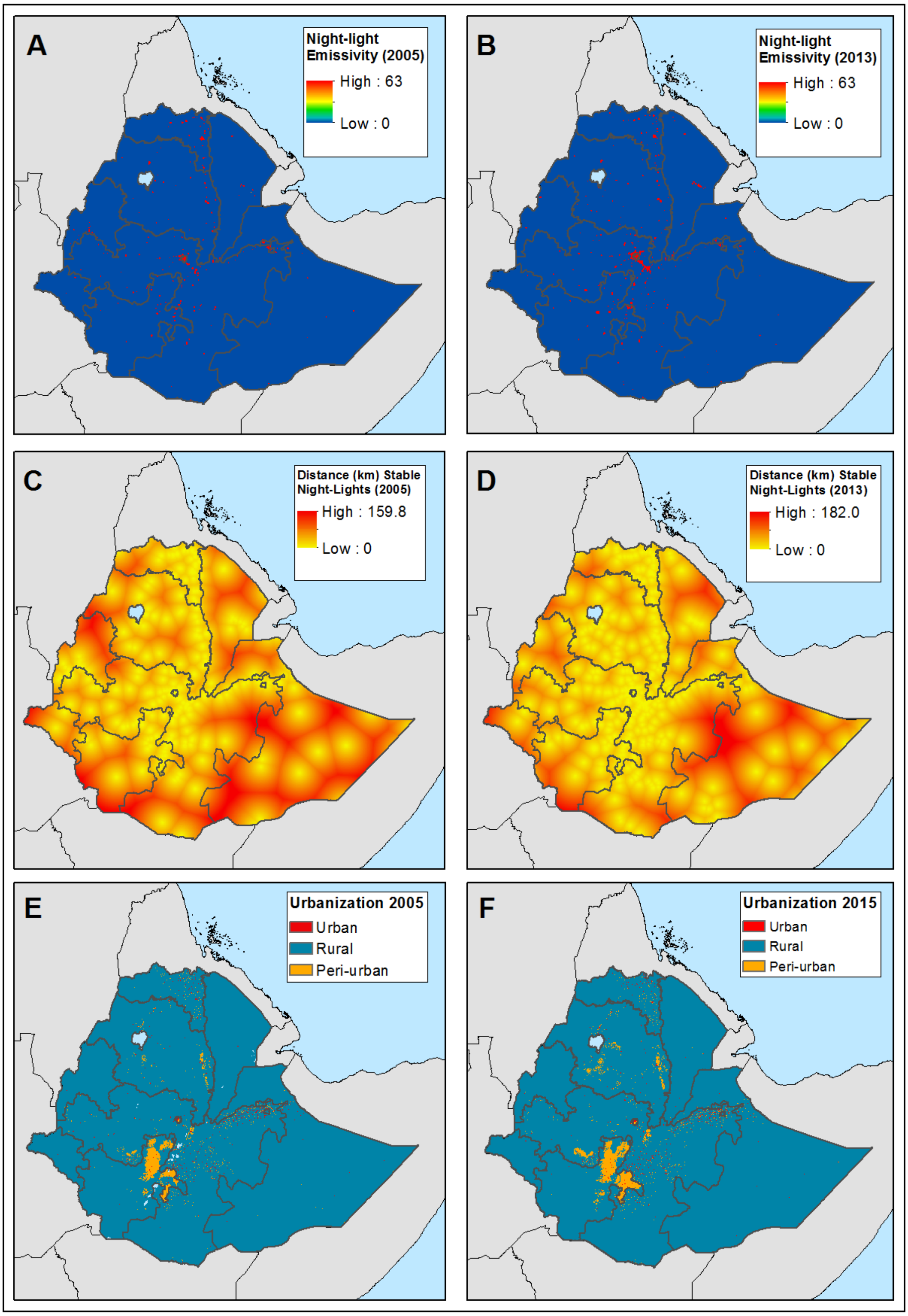

**Figure S8** Empirical semivariogram of log-transformed of untreated obstetric fistula in 2005 (A) and 2016 (B). Shaded blue area corresponds to confidence intervals for random spatial patterns obtained via simulation.

**A**

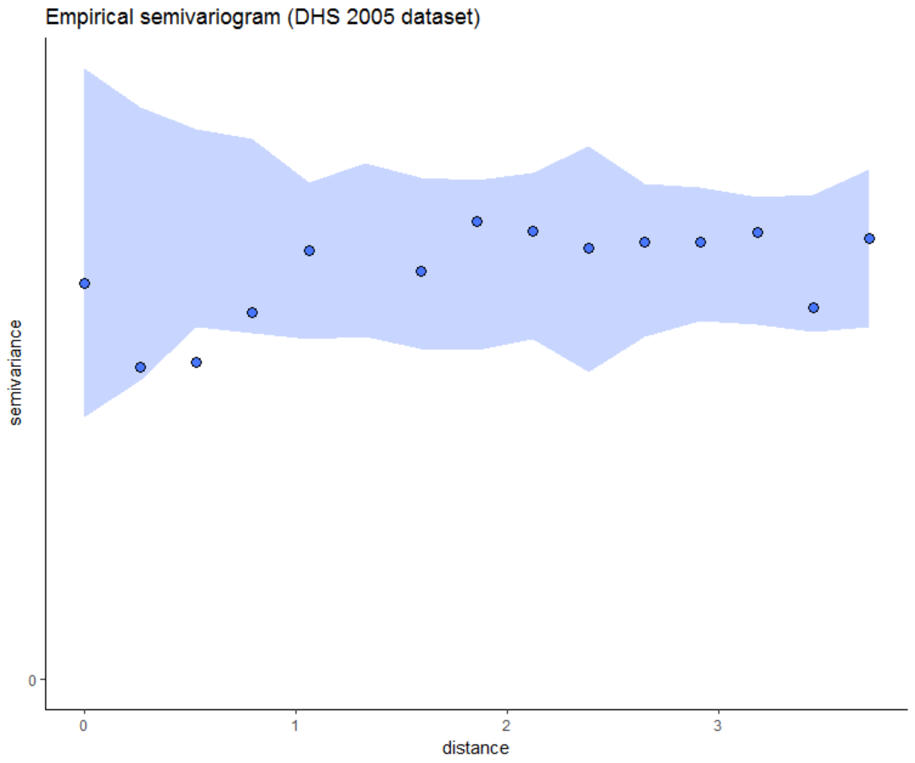

**B**

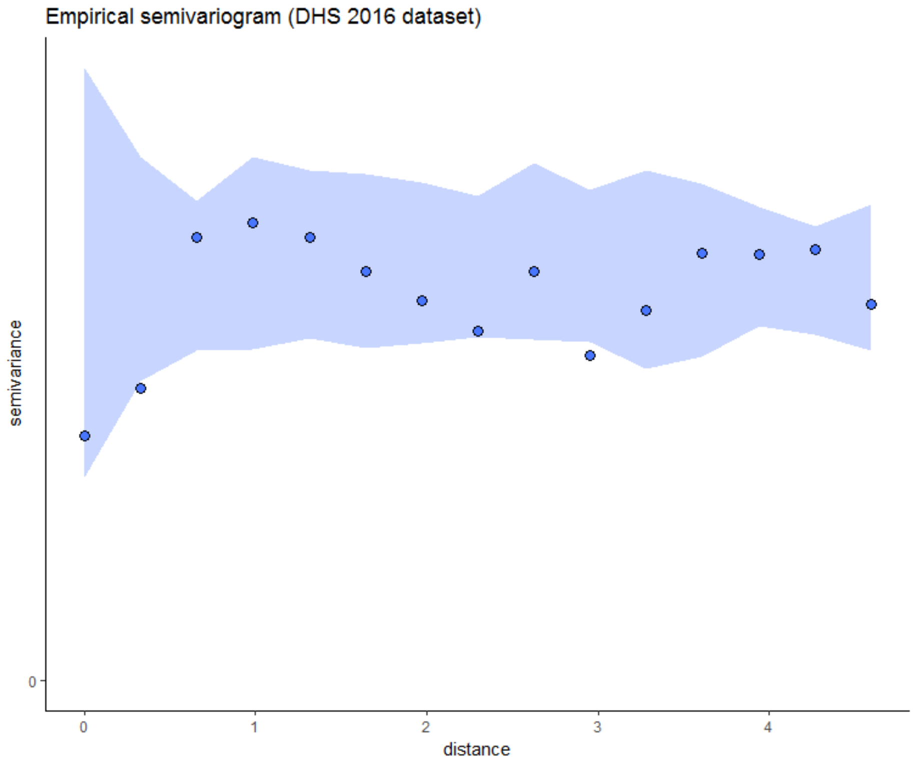

**Figure S9** Tests via simulation to determine the predictive performance of binomial mixed models of lifetime (1) and untreated obstetric fistula (2) (Ethiopia, 2005) fitted with the following covariates: i) Accessibility (min) to major towns in 2000, ii) proportion women reporting four antenatal visits, iii) proportion literate women, and iv) Cost distance to nearest health facility.

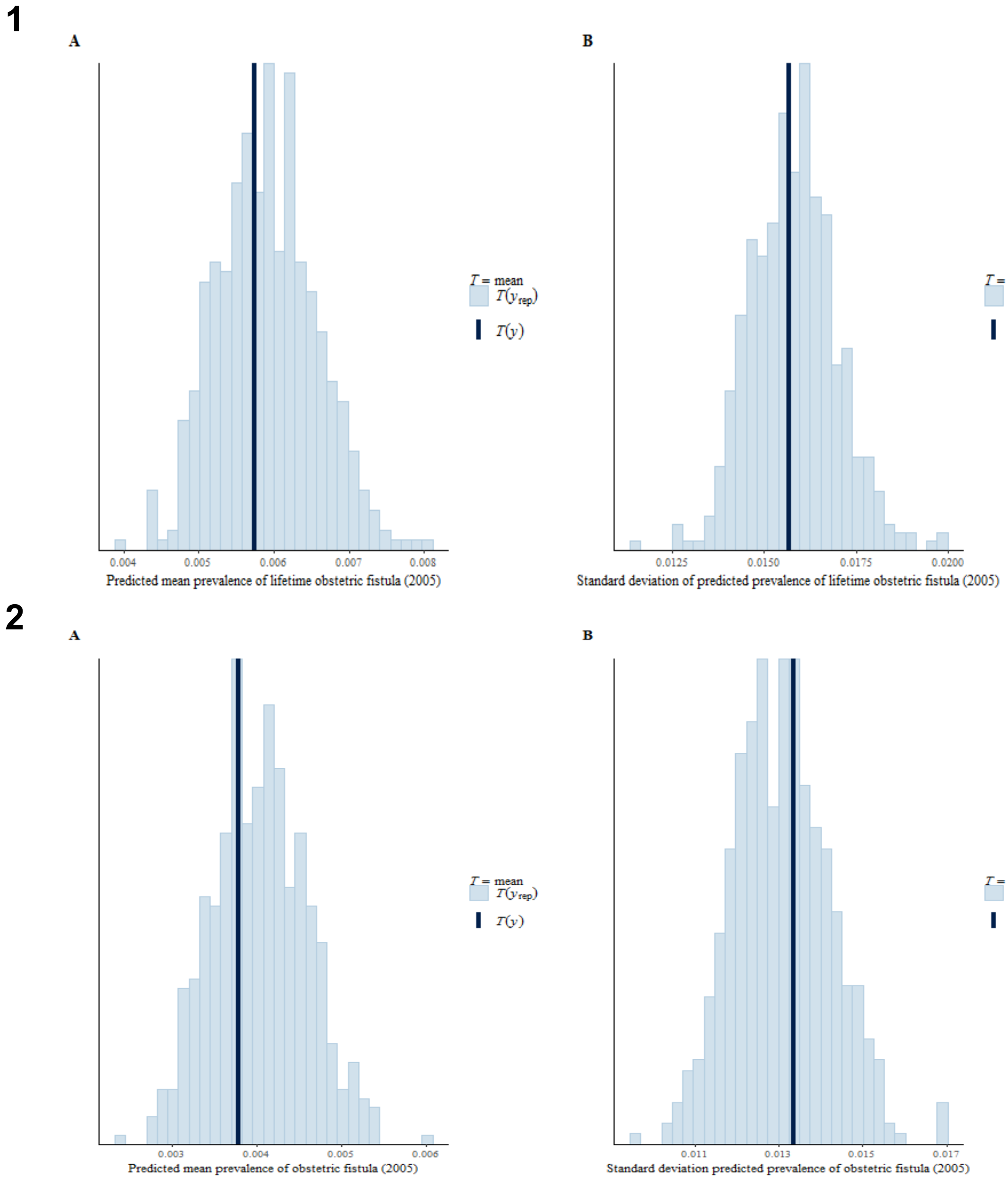

**Figure S10** Tests via simulation to determine the predictive performance of binomial mixed models of lifetime (1) and untreated obstetric fistula (2) (Ethiopia, 2016) fitted with the following covariates: i) Accessibility (min) to major towns in 2015, ii) proportion women having delivered in a health facility, iii) proportion women reporting using modern contraception methods, and iv) Cost distance to nearest health facility.

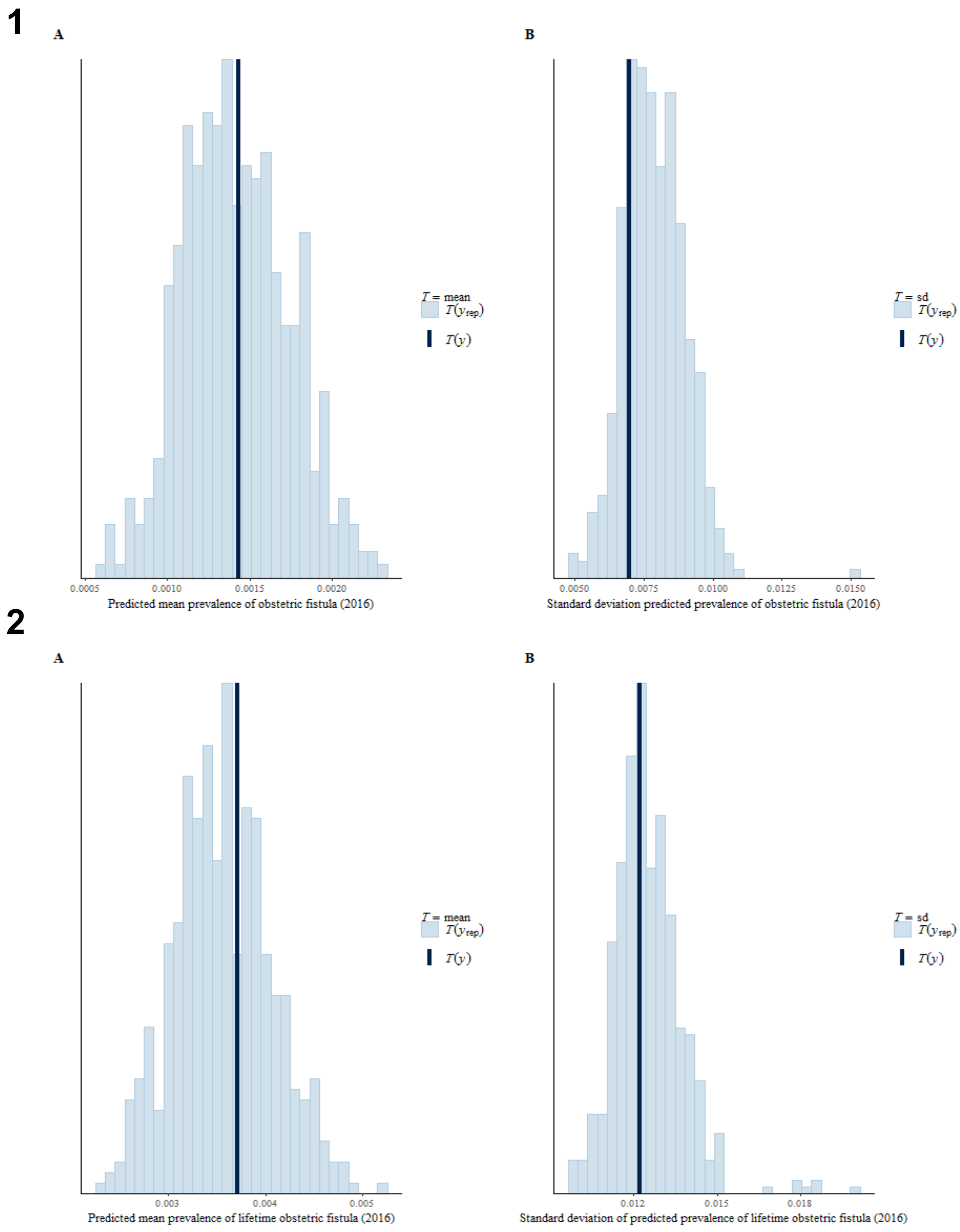

**Figure S11** Predicted mean prevalence of lifetime obstetric fistula in 2005 (A) and 2016 (B) across Ethiopia. Prevalence is provided in cases per 1,000 women of childbearing age.

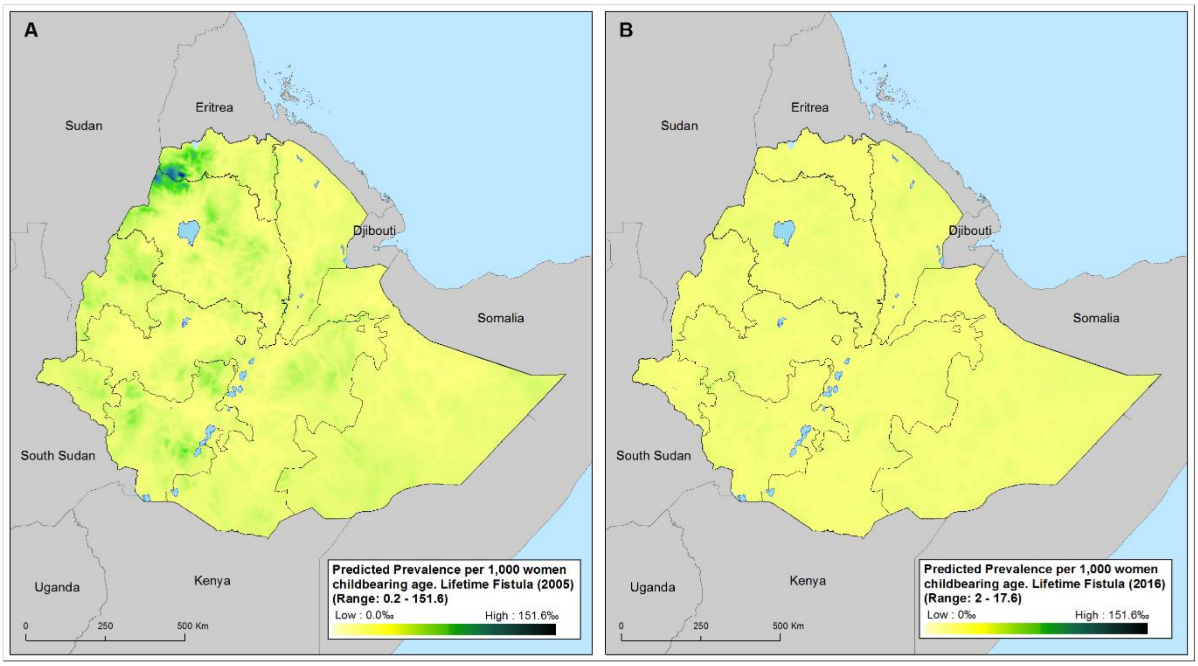

**Figure S12** Estimated number of childbearing women (15-49 year-old) suffering from lifetime obstetric fistula in 2005 (A) and in 2016 (B). Figures have been obtained using the estimated number of females aged between 15 and 49 per sq-km estimated by the WorldPop project.

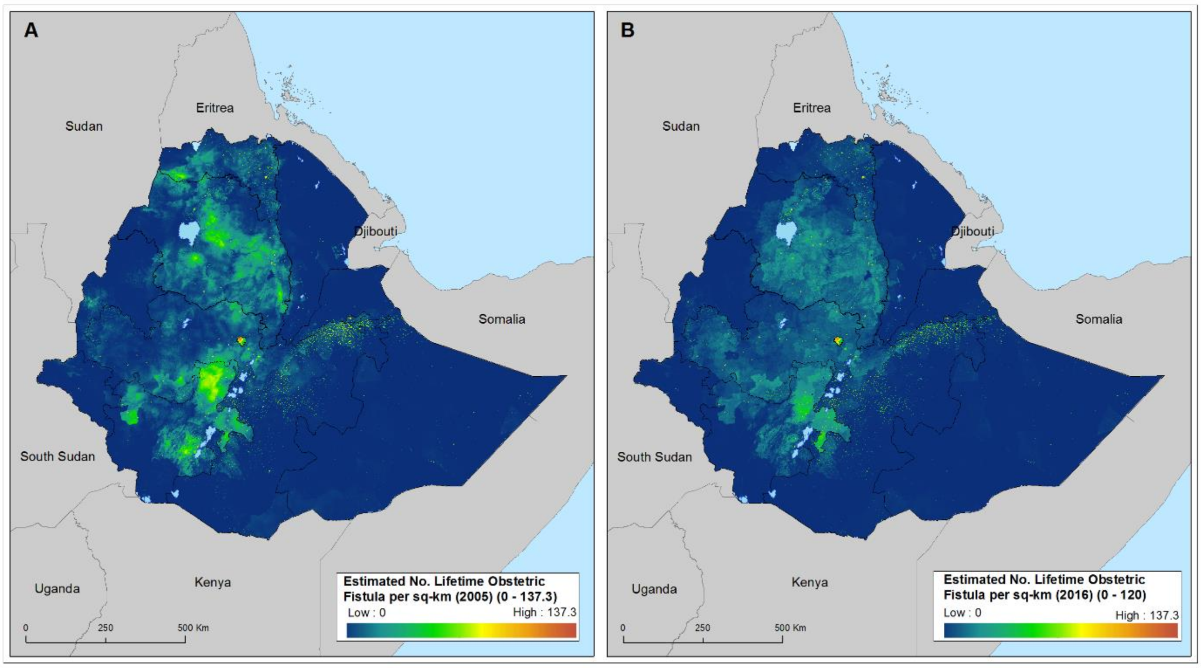

**Figure S13** Predicted mean prevalence of lifetime obstetric fistula by district in 2005 (A) and 2016 (B) across Ethiopia. Prevalence is provided in cases per 1,000 women of childbearing age.

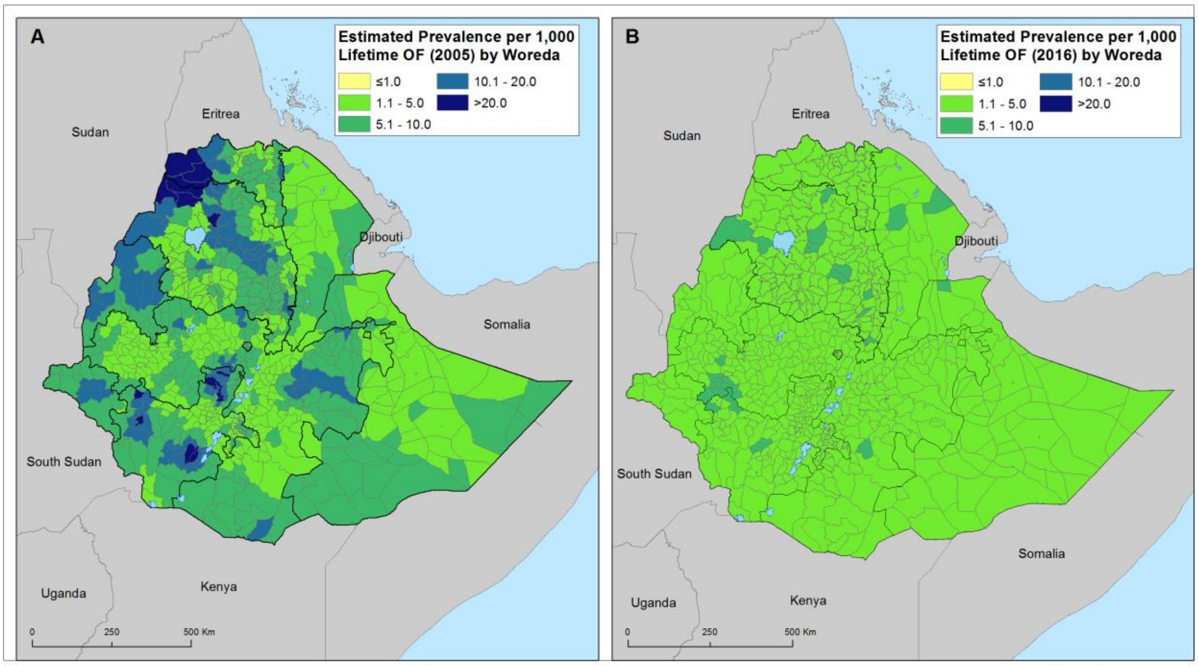

**Figure S14** Estimated number of childbearing women (15-49 year-old) suffering from lifetime obstetric fistula by district in 2005 (A) and in 2016 (B). Figures have been obtained using the estimated number of females aged between 15 and 49 per sq-km estimated by the WorldPop project.

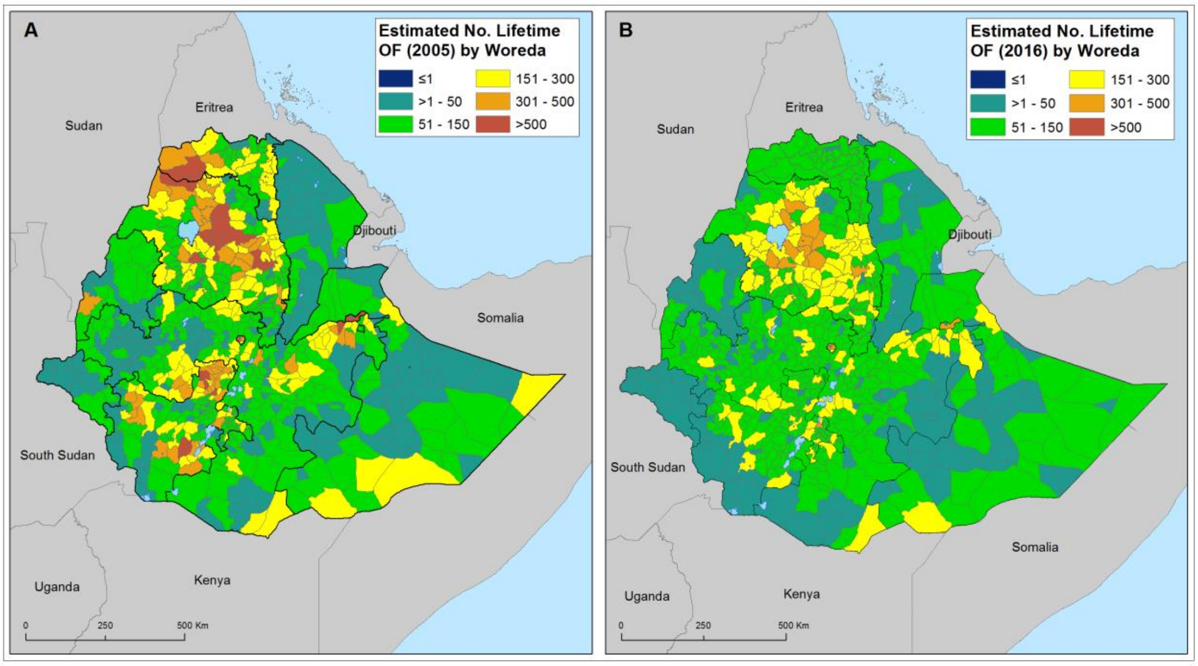

**Table S1.** Correlation matrix of shortlisted covariates (continuous data) to be used in the modelling of untreated and lifetime obstetric fistula in 2005 in Ethiopia. Spearman's correlation test was used to explore the collinearity between pairs of covariates.

| | **Accessibility 2000** | ***Prop. Attending Antenatal Visits** | ***Prop. Assisted Deliveries** | **Cost Dist. Nearest HF** | ***Prop. Literate Women** | ***Prop. Using Contraception** | **Night-Light Emissivity 2005** | **Dist. Stable Nigh-lights 2005** |
|---|---|---|---|---|---|---|---|---|
| **Accessibility 2000** | 1.000 | | | | | | | |
| ***Prop. Attending Antenatal Visits** | -0.152 | 1.000 | | | | | | |
| ***Prop. Assisted Deliveries** | -0.228 | 0.654 | 1.000 | | | | | |
| **Cost Dist. Nearest HF** | 0.341 | -0.406 | -0.368 | 1.000 | | | | |
| ***Prop. Literate Women** | -0.223 | 0.617 | 0.568 | -0.452 | 1.000 | | | |
| ***Prop. Using Contraception** | -0.126 | 0.679 | 0.580 | -0.439 | 0.721 | 1.000 | | |
| **Night-Light Emissivity 2005** | -0.063 | 0.127 | 0.151 | -0.055 | 0.139 | 0.083 | 1.000 | |
| **Dist. Stable Nigh-lights 2005** | 0.302 | -0.396 | -0.351 | 0.524 | -0.492 | -0.451 | -0.076 | 1.000 |

*Estimates for 2016

**Table S2.** Correlation matrix of shortlisted covariates (continuous data) to be used in the modelling of untreated and lifetime obstetric fistula in 2016 in Ethiopia. Spearman's correlation test was used to explore the collinearity between pairs of covariates.

| | **Accessibility 2015** | ***Prop. Attending Antenatal Visits** | ***Prop. Assisted Deliveries** | **Cost Dist. Nearest HF** | ***Prop. Literate Women** | ***Prop. Using Contraception** | **Night-Light Emissivity 2013** | **Dist. Stable Nigh-lights 2013** |
|---|---|---|---|---|---|---|---|---|
| **Accessibility 2016** | 1.000 | | | | | | | |
| ***Prop. Attending Antenatal Visits** | -0.381 | 1.000 | | | | | | |
| ***Prop. Assisted Deliveries** | -0.335 | 0.654 | 1.000 | | | | | |
| **Cost Dist. Nearest HF** | 0.765 | -0.406 | -0.368 | 1.000 | | | | |
| ***Prop. Literate Women** | -0.415 | 0.617 | 0.568 | -0.452 | 1.000 | | | |
| ***Prop. Using Contraception** | -0.401 | 0.679 | 0.580 | -0.439 | 0.721 | 1.000 | | |
| **Night-Light Emissivity 2013** | -0.082 | 0.148 | 0.186 | -0.072 | 0.168 | 0.106 | 1.000 | |
| **Dist. Stable Nigh-lights 2013** | 0.523 | -0.382 | -0.330 | 0.480 | -0.448 | -0.452 | -0.092 | 1.000 |

*Estimates for 2016

**Table S3.** Relationship between risk of untreated and lifetime obstetric fistula in Ethiopia in 2005 and each potential explanatory variable.

Variables were grouped according to its nature and collinearity. Selection of variables was made by fitting univariate models relating the logit of obstetric fistula to each of the covariate and eventually comparing the models in terms of the Akaike Information Criterion (AIC), choosing those variables which yielded the lowest AIC value and allowed for better fitting in multivariate binomial model. Highlighted in grey the covariates selected to fit the binomial regression models.

| Predictors | AIC value | |
|---|---|---|
| | Current OF | Life time OF |
| Accessibility 2000 | 428.51 | 562.43 |
| Urbanization 2005(Rural/ Urban) | 430.94 | 564.13 |
| Nightlight Emissivity 2005 | 429.08 | 563.22 |
| Distance to stable night light 2005 | 430.17 | 562.63 |
| Cost distance to nearest health facility | 435.23 | 562.96 |
| *Proportion of assisted deliveries | 435.13 | 564.28 |
| *Proportion of using contraception | 434.24 | 564.29 |
| *Proportion of literate women | 434.24 | 563.95 |

OF: Obstetric Fistula / * Based on DHS 2016 data

**Table S4.** Relationship between risk of untreated and lifetime obstetric fistula in Ethiopia in 2016 and each potential explanatory variable.

Variables were grouped according to its nature and collinearity. Selection of variables was made by fitting univariate models relating the logit of obstetric fistula to each of the covariate and eventually comparing the models in terms of the Akaike Information Criterion (AIC), choosing those variables which yielded the lowest AIC value and allowed for better fitting in multivariate binomial model. Highlighted in grey the covariates selected to fit the binomial regression models.

| **Predictors** | **AIC value** | |
|---|---|---|
| | **Current OF** | **Life time OF** |
| Accessibility 2016 | 229.61 | 446.24 |
| Urbanization 2015(Rural/ Urban) | 230.84 | 444.57 |
| Nightlight Emissivity 2013 | 231.11 | 446.25 |
| Distance to stable night light 2013 | 233.04 | 448.23 |
| Cost distance to nearest health facility | 229.56 | 446.56 |
| *Proportion of women attending antenatal visits | 233.07 | 448.10 |
| *Proportion of assisted deliveries | 232.52 | 447.67 |
| *Proportion of using contraception | 231.44 | 448.06 |
| *Proportion of literate women | 233.21 | 448.34 |

OF: Obstetric Fistula / * Based on DHS 2016 data

**Table S5** Regression coefficients and credible intervals for the covariates used to fit the multivariate binomial mixed models of untreated and lifetime obstetric fistula in 2005

| | Untreated Fistula 2005 | | | Lifetime Fistula 2005 | | |
|---|---|---|---|---|---|---|
| | Coeffs | 95%CI | Pr(>\|z\|) | Coeffs | 95%CI | Pr(>\|z\|) |
| (Intercept) | 0.004 | (0.003-0.006) | < 2e-16 | 0.006 | (0.005-0.008) | < 2e-16 |
| Accessibility 2000 | 1.536 | (0.908-2.630) | 0.113 | 1.630 | (1.064-2.533) | 0.028 |
| Accessibility 2000 - QT | 0.774 | (0.556-1.001) | 0.090 | 0.843 | (0.679-1.004) | 0.094 |
| Cost Dist. Nearest HF | 0.960 | (0.679-1.277) | 0.800 | 0.936 | (0.700-1.195) | 0.624 |
| % Women Attending 4 Antenatal Visits | 1.902 | (1.196-3.027) | 0.007 | 2.021 | (1.394-2.925) | 0.000 |
| % Literate Women | 0.529 | (0.320-0.868) | 0.012 | 0.598 | (0.399-0.891) | 0.012 |

QT: Quadratic term; HF: Health Facility

**Table S6** Regression coefficients and credible intervals for the covariates used to fit the multivariate binomial mixed models of untreated and lifetime obstetric fistula in 2016

| | Untreated Fistula 2016 | | | Lifetime Fistula 2016 | | |
|---|---|---|---|---|---|---|
| | Coeffs | 95%CI | Pr(>\|z\|) | Coeffs | 95%CI | Pr(>\|z\|) |
| (Intercept) | 0.001 | (0.001-0.002) | <2e-16 | 0.003 | (0.003-0.004) | <2e-16 |
| Accessibility 2015 | 1.252 | (0.676-2.456) | 0.479 | 1.169 | (0.747-1.879) | 0.500 |
| Cost Dist. Nearest HF | 1.236 | (0.578-1.993) | 0.482 | 1.042 | (0.605-1.518) | 0.859 |
| % Women Delivering at HF | 0.750 | (0.421-1.316) | 0.318 | 0.878 | (0.611-1.257) | 0.477 |
| % Women Using Contraception | 1.794 | (1.067-3.044) | 0.028 | 1.218 | (0.872-1.697) | 0.244 |

HF: Health Facility
